# Local circuit allowing hypothalamic control of hippocampal area CA2 activity and consequences for CA1

**DOI:** 10.1101/2020.09.18.303693

**Authors:** Vincent Robert, Ludivine Therreau, Arthur J.Y. Huang, Roman Boehringer, Denis Polygalov, Thomas McHugh, Vivien Chevaleyre, Rebecca A. Piskorowski

**Affiliations:** Université de Paris, INSERM UMR1266, Institute of Psychiatry and Neuroscience of Paris, Team Synaptic Plasticity and Neural Networks, 102-108 rue de la santé, 75014, Paris, France; GHU PARIS Psychiatrie and Neurosciences, F-75014 Paris, France; Laboratory for Circuit and Behavioral Physiology, RIKEN Center for Brain Science, 2-1 Hirosawa, Wakoshi, Saitama, Japan

## Abstract

The hippocampus is critical for memory formation. The hypothalamic supramammillary nucleus (SuM) sends long-range projections to hippocampal area CA2. While the SuM-CA2 connection is critical for social memory, how this input acts on the local circuit is unknown. We found that SuM axon stimulation elicited mixed excitatory and inhibitory responses in area CA2 pyramidal neurons (PNs). We found that parvalbumin-expressing basket cells as responsible for the feedforward inhibitory drive of SuM over area CA2. Inhibition recruited by the SuM input onto CA2 PNs increased the precision of action potential firing both in conditions of low and high cholinergic tone. Furthermore, SuM stimulation in area CA2 modulates CA1 activity, indicating that synchronized CA2 output drives a pulsed inhibition in area CA1. Hence, the network revealed here lays basis for understanding how SuM activity directly acts on the local hippocampal circuit to allow social memory encoding.

## Introduction

The hippocampus is critical for memory formation and spatial navigation (Buzsáki and Moser, 2013; Eichenbaum and Cohen, 2014), yet basic questions persist regarding the circuitry and cellular components allowing these processes. While area CA2 has been shown to play a significant role in several hippocampal processes including social memory formation (Hitti and Siegelbaum, 2014; Stevenson and Caldwell, 2014) sharp-wave ripple generation (Oliva et al., 2016) and spatial encoding (Kay et al., 2016), information about the local circuitry and cellular processes allowing these functions is lacking. There is mounting evidence that generalizations cannot be made from the rich understanding of areas CA1 and CA3, as neurons in area CA2 have been shown to have unique molecular expression profiles (Cembrowski et al., 2016; Lein et al., 2004), morphology (Bartesaghi and Ravasi, 1999; No, 1934) and cellular properties (Robert et al., 2020; Srinivas et al., 2017; Sun et al., 2014). Notably, and in contrast to area CA1, CA2 pyramidal neurons do not undergo NMDA-mediated synaptic plasticity (Dasgupta et al., 2020; Zhao et al., 2007). Rather, the excitability of this region is tightly controlled by a highly plastic network of inhibitory neurons (Leroy et al., 2017; Nasrallah et al., 2015; Piskorowski and Chevaleyre, 2013). When active, CA2 pyramidal neurons (PNs) can strongly drive area CA1 (Chevaleyre and Siegelbaum, 2010; Kohara et al., 2014; Nasrallah et al., 2019), thereby influencing hippocampal output. Furthermore, CA2 neurons also project to area CA3, where they recruit inhibition (Boehringer et al., 2017; Kohara et al., 2014) and act to control hippocampal excitability. Thus, CA2 neurons are poised to have long-reaching effects in the hippocampus, and a better understanding of the regulation of this region is needed.

The hypothalamic supramammillary (SuM) nucleus sends projections to both area CA2 and the dentate gyrus (Haglund et al., 1984; Vertes, 1992). These long-range connections have been shown in several species including rodents, primates and humans (Berger et al., 2001; Haglund et al., 1984; Wyss et al., 1979) where they are present in early hippocampal development. The SuM has been found to be active during a wide variety of conditions including novel environment exposure (Ito et al., 2009), reinforcement learning (Ikemoto, 2005; Ikemoto et al., 2004), food anticipation (May et al., 2019), and during REM sleep and arousal (Pedersen et al., 2017; Renouard et al., 2015). This nucleus is also known for participating in hippocampal theta rhythm (Pan and McNaughton, 2002, 1997), possibly by its direct projection to the hippocampus or by modulation of the medial septum (Borhegyi et al., 1998; Vertes and Kocsis, 1997) and regulating spike-timing between hippocampus and the cortex (Ito et al., 2018). Disruption of SuM neuron activity with pharmacological methods (Aranda et al., 2008; Shahidi et al., 2004) or lesions (Aranda et al., 2006) has been reported to disrupt hippocampal memory. Serotonin depletion of the SuM leads to deficiencies in spatial learning in the Morris water maze, and results in altered hippocampal theta activity (Gutiérrez-Guzmán et al., 2012; Hernández-Pérez et al., 2015). Salient rewarding experiences also activate the SuM, as evidenced by cFos expression in monoaminergic SuM neurons by consumption of rewarding food (Plaisier et al., 2020). Furthermore, the rewarding aspects of social aggression have been shown to involve an excitatory circuit between the hypothalamic ventral premammillary nucleus and the SuM (Stagkourakis et al., 2018). It has recently been shown that there are two separate populations of cells in the SuM that target either CA2 or the DG (Chen et al., 2020). In the DG, the SuM terminals release both glutamate and GABA (Boulland et al., 2009; Hashimotodani et al., 2018; Pedersen et al., 2017; Soussi et al., 2010). The SuM-DG projection has been recently shown to play a role in modulating DG activity in response to contextual novelty (Chen et al., 2020) and spatial memory retrieval (Li et al., 2020). In contrast, functional studies of the SuM-CA2 projection have found that this connection is entirely glutamatergic (Chen et al., 2020). It was recently discovered that the CA2-projecting SuM neurons are active during social novelty exposure, and their selective stimulation prevents expression of a memory of a familiar conspecific (Chen et al., 2020). These findings strongly suggest that the SuM-CA2 connection conveys a social novelty signal to the hippocampus. Furthermore, recent in vivo recordings from the SuM in anaesthetized rats recently reported that a subset of SuM neurons were active earlier than CA2 and other hippocampal cells during SWR (Vicente et al., 2020), indicating a possible role for the SuM-CA2 projection in shaping area CA2 activity prior to SWR onset.

Even with the anatomical and in vivo data, the properties and consequences of SuM activation on area CA2 activity remain unexplored. In this study, we use a combination of approaches to specifically examine the effects of SuM input stimulation on neuronal activity in hippocampal area CA2. Here, we show that the SuM-evoked post-synaptic excitation of CA2 PN is controlled by SuM-driven inhibition. We identified PV-expressing basket cells as the neuronal population most strongly excited by SuM input in area CA2, and thus likely responsible for the feedforward inhibition evoked by SuM in CA2 PNs. We found that recruitment of this inhibition enhances the precision of AP firing by area CA2 PNs in conditions of low and high cholinergic tone. Finally, we observed that the resulting synchronized CA2 PN activity drives inhibition in area CA1, thereby providing a circuit mechanism through which SuM can modulate hippocampal excitability by controlling area CA2 output.

## Results

In order to functionally investigate the SuM projection to area CA2, we used an anterograde strategy in two separate transgenic mouse lines (Figure 1A and Supplemental Figure 1F). It has been shown that the source of vesicular glutamate transporter 2 (VGluT2)-immunopositive boutons in area CA2 originate from the SuM (Halasy et al., 2004). To further assess where these VGluT2-expressing SuM cells project into the hippocampus, we injected an AAV to express channelrhodopsin(H143R)-YFP (ChR2-EYFP) under the control of Cre into the SuM of a transgenic mouse line with Cre expression controlled by the VGluT2 promoter, the Tg(Slc17ab-icre)10Ki line (Borgius et al., 2010) (Supplemental Figure 1F). In parallel, we used a novel mouse line, the Csf2rb2-Cre line that selectively expresses Cre in the SuM (Chen et al., 2020) (Figure 1A). To find the optimal injection site, we injected a retrograde canine adenovirus type 2 (CAV-2) into area CA2 of the hippocampus to permit the expression of Cre-recombinase (Cre) in hippocampal-projecting SuM neurons, and an adeno-associated virus (AAV) was injected into the SuM to allow the expression of EGFP under the control of Cre (Supplemental Figure 1A). In 5 animals the injection of retrograde CAV-2 was sufficiently targeted to area CA2, as indicated by the presence of EGFP-expressing SuM axonal fibers primarily in this hippocampal area (Supplemental Figure 1B). We stained for calretinin to define the boundaries of the SuM nucleus (Pan and Mcnaughton, 2004). Consistent with what has been described (Chen et al., 2020), we observed that CA2-projecting cells co-express calretinin and are located in the medial SuM (Supplemental figure 1C-D). These cells were located bilaterally, ventral to the fiber bundles that traverse the SuM (Supplemental Figure 1C).

**Figure 1.**
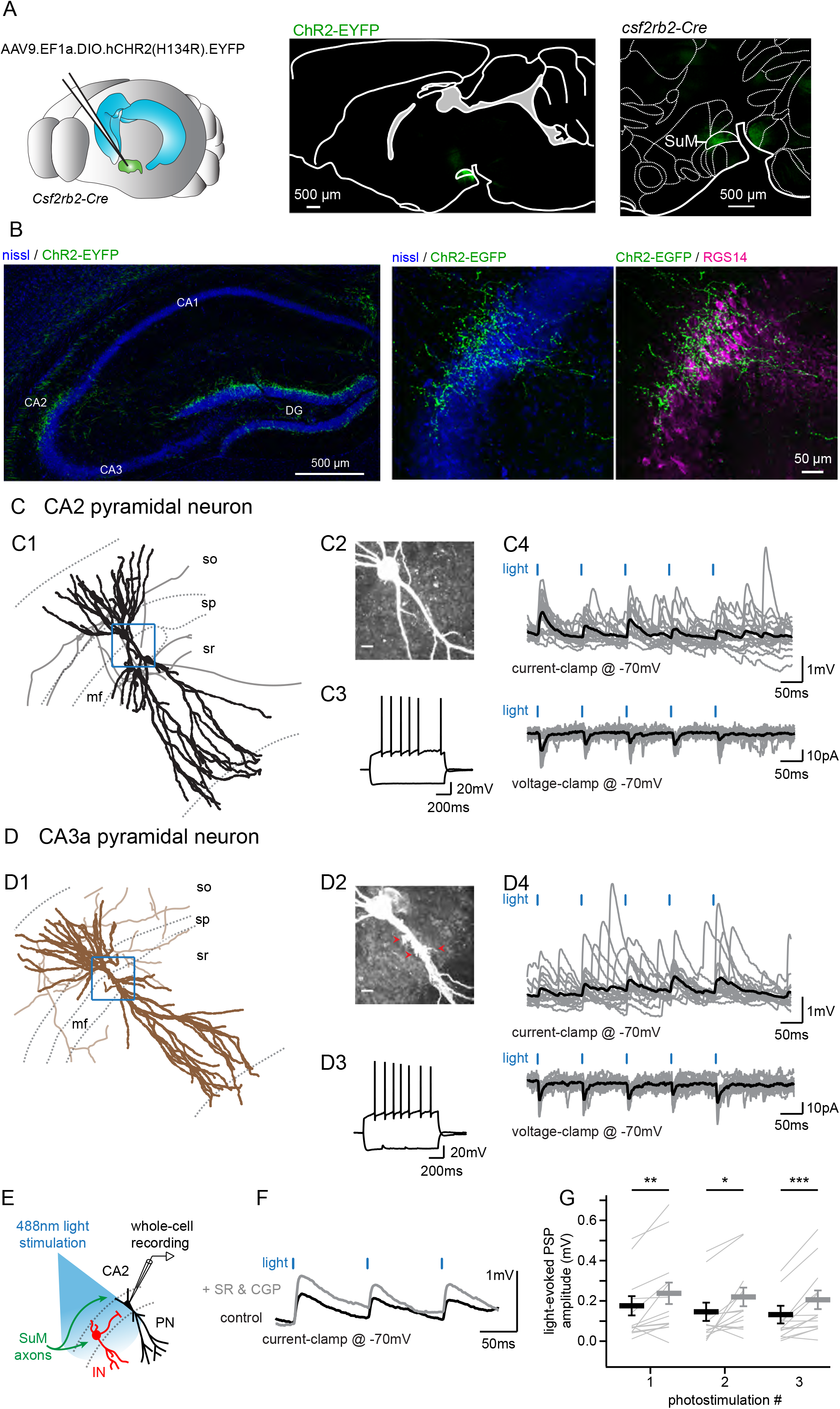
Selective functional mapping of SuM neurons that project to hippocampal area CA2. A. Left, diagram illustrating the injection of AAVs into the SuM. Middle, sagittal image indicating the infected SuM area expressing hCHR2(H134R)-EYFP (green). Right, expanded view of injection site in the Csf2rbr-Cre mouse line. B. Left, hCHR2(H134R)- EYFP -expressing SuM fibers (green) and nissl staining (blue) in the hippocampus. Right, higher magnification image of area CA2 with hCHR2(H134R)-EYFP -expressing SuM fibers (green) and nissl staining (blue) and RGS14 staining (magenta) to label area CA2. C. CA2 pyramidal neurons in the SuM-innervated region receive excitatory transmission. (C1) Example CA2 PN reconstruction (dendrites in black, axons in grey, hippocampal stratum borders shown in dotted line, area demarcated in blue corresponds to the expanded image in C2). (C2) Biocytin labeling of the recorded cell proximal dendrites, scale bar represents 10 μm. (C3) AP firing and repolarizing sag current in response to steps of +800 and −400 pA current injection. (C4) Light-evoked EPSPs (top traces, individual traces shown in grey, average trace shown in black) and EPSCs (bottom traces, individual traces shown in grey, average trace shown in black). **D.** CA3 pyramidal neurons in the SuM-innervated region receive excitatory transmission. (D1) Example CA3 PN reconstruction (dendrites in brown, axons in light brown, hippocampal stratum borders shown in dotted line, area demarcated in blue corresponds to the expanded image in D2). (D2) Biocytin labeling of the recorded cell proximal dendrites, note the presence of thorny excrescences, as indicated by the red arrows; scale bar represents 10 μm. (D3) AP firing and repolarizing sag current in response to steps of +800 and −400 pA current injection. (D4) Light-evoked EPSPs (top traces, individual traces shown in grey, average trace shown in black) and EPSCs (bottom traces, individual traces shown in grey, average trace shown in black). E. Diagram illustrating the whole-cell recordings of area CA2 PNs and SuM fiber light stimulation in acute slice preparation. F. Sample traces of three 10 Hz SuM light-evoked PSPs before and after blocking inhibitory transmission (control shown in black, SR95531 & CGP55845A shown in grey). G. Summary graph of light-evoked PSP amplitudes recorded in PNs before and after application of 1 μM SR95531& 2 μM CGP55845A (individual cells shown as thin lines, population average shown as thick line, error bars represent SEM, n = 14; Wilcoxon signed-rank tests, p = 0.004 for the first PSP, p = 0.013 for the second PSP, p < 0.001 for the third PSP).

We found that with both transgenic mouse lines we could reproducibly restrict expression of ChR2-EYFP in the SuM and avoid infecting nearby hypothalamic regions that also may project to the hippocampus (Figure 1A, Supplemental Figure 1F). For all experiments, injection sites were examined post hoc to ensure correct targeting of the SuM. With both lines of transgenic mice, we observed identical patterns of SuM fiber localization in the hippocampus. EYFP-containing SuM axons were found throughout the supragranular layer of the DG and in area CA2 (Figure 1B) where they clustered around the pyramidal layer (stratum pyramidale, SP) and spread in stratum oriens (SO). The SuM fiber projection area was clearly restricted to area CA2, as defined by expression of the CA2-specific markers PCP4 and RGS14 and did not spread to neighboring areas CA3 and CA1 (Figure 1B). In order to maximize the precision of our experiments, we frequently only achieved partial infection of the SuM, as indicated by the sparseness of ChR2-EYFP-containing fibers in comparison to the number of vGluT2-stained boutons in this region (Supplemental Figure 1G-H).

### SuM axons provide excitatory glutamatergic input to pyramidal neurons in area CA2 and CA3a

In order to better understand the cellular targets and consequences of SuM input activity in area CA2, we applied the above experimental strategy to express ChR2-EYFP in SuM axonal fibers and performed whole-cell current and voltage clamp recordings of PNs across the hippocampal CA regions and activated projecting axons with pulses of 488 nm light in acute hippocampal slices. Following all recordings, we performed post-hoc anatomical reconstructions of recorded cells and axonal fibers, as well as immunohistochemical staining for CA2-area markers.

We observed that photostimulation of SuM axons elicited excitatory post-synaptic responses in 63 % of PNs (n = 166 of 263 cells) located in area CA2. PNs in this region shared similar overall dendritic morphologies and electrophysiological properties (Table 1) but differed along two criteria. First, in *stratum lucidum* some PNs clearly had thorny excrescences (TE) while others had very smooth apical dendrites (Figure 1C-D). Based on the presence of TEs, we classified cells as CA2 or CA3 PNs (unequivocal distinction was possible for 148 neurons). Second, the distribution locations of PN soma along the radial axis of the hippocampus allowed us to cluster them as deep (closer to *stratum oriens*, SO) or superficial (closer to *stratum radiatum*, SR) subpopulations (unequivocal distinction was possible for 157 neurons). We found that the SuM-PN connectivity was not different between CA2 and CA3 PNs (Table 2, χ^2^ test for CA2 and CA3 PNs, p = 0.572) or between deep and superficial PNs (Table 2, χ^2^ test for deep and superficial PNs, p = 0.946). Light-evoked excitatory post-synaptic potentials (EPSPs) and excitatory post-synaptic currents (EPSCs) recorded at −70mV were of fairly small amplitude (Figure 1C and 1D) that were similar regardless of the PN type or somatic location (Table 2, Mann-Whitney U test for CA2 and CA3 PNs, p = 0.409; Mann-Whitney U test for deep and superficial PNs, p = 0.306). Because no significant differences in post-synaptic responses to SuM input stimulation were observed between CA2 and CA3 PNs as well as between deep and superficial PNs, data from all PNs was pooled for the rest of the study. The small amplitude of SuM input-evoked post-synaptic responses in PNs was not due to under-stimulation of SuM axons as EPSC amplitudes rapidly reached a plateau when increasing light intensity (Supplemental Figure 2). We are confident that this transmission is due to action potential-generated vesicle release because all transmission was blocked following application of tetrodotoxin (TTX) (Supplemental Figure 2). The pure glutamatergic nature of the SuM input was confirmed by the complete block of light-evoked synaptic transmission following the application of NBQX and D-APV (Supplemental Figure 2; amplitudes were 16 ± 4.8 pA in control and 1.8 ± 0.3 pA in NBQX & D-APV, n = 6; Wilcoxon signed-rank test, p = 0.03). These data confirm that SuM inputs provide long-range glutamatergic excitation to CA2 and CA3 PNs in area CA2.

**Table 1.**
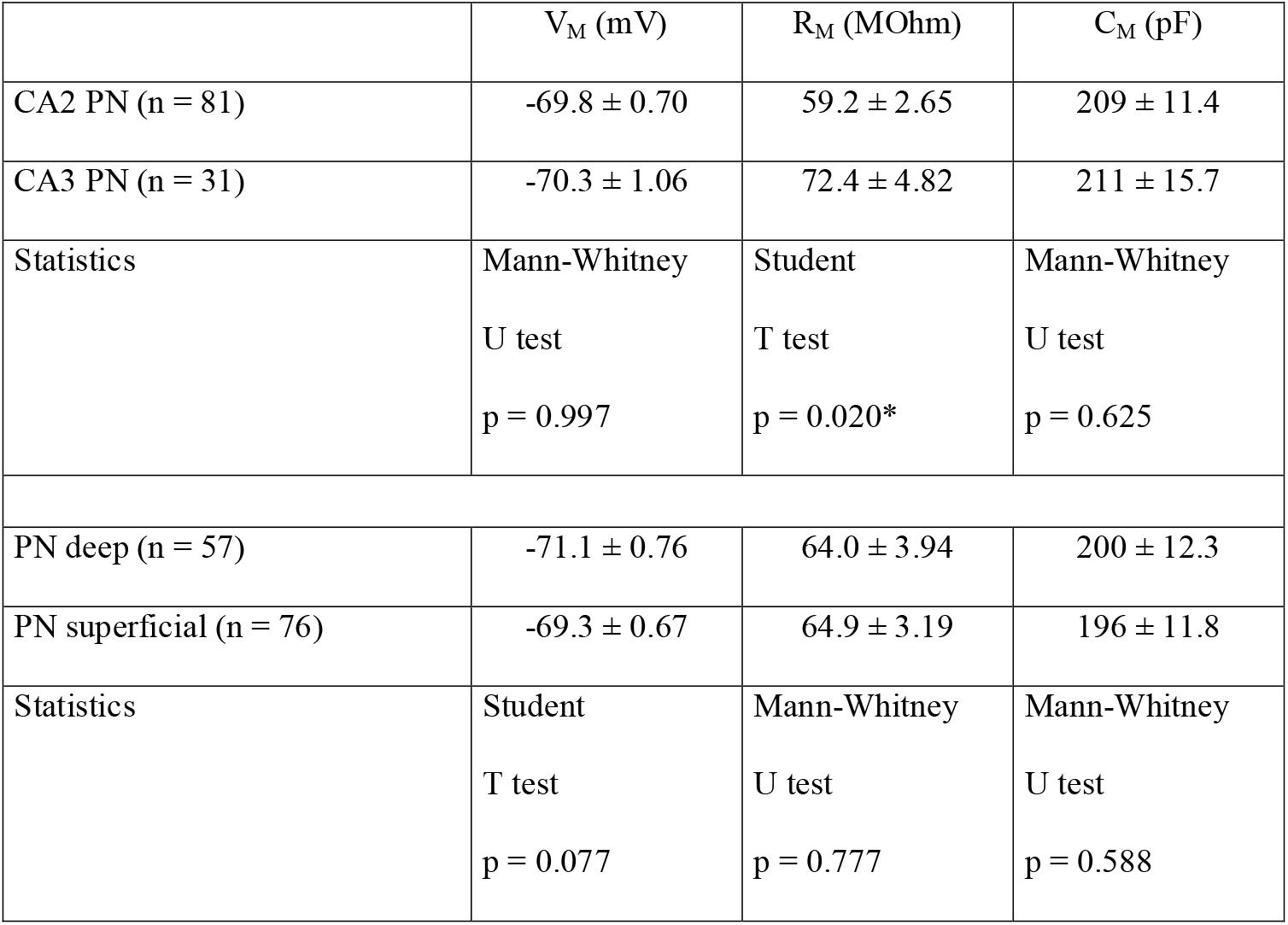
Electrophysiological properties of pyramidal neurons in SuM-innervated area.

**Table 2.** Characteristics of SuM light-evoked transmission onto pyramidal neurons.

|  | EPSC |  |  |  |  |  |
| --- | --- | --- | --- | --- | --- | --- |
| cell type | connectivity (%) | amplitude (pA) | rise time (ms) | decay time (ms) | latency (ms) | success rate |
| CA2 PN | 56 (n = 58 of 103) | $16 \pm 1.9$ | $2.9 \pm 0.1$ | $14 \pm 0.8$ | $2.4 \pm 0.2$ | $0.44 \pm 0.03$ |
| CA3 PN | 49 (n = 22 of 45) | $23 \pm 5.9$ | $3.0 \pm 0.2$ | $14 \pm 0.9$ | $2.7 \pm 0.3$ | $0.56 \pm 0.06$ |
| Statistics | $\chi^2$ test<br>p = 0.572 | Mann-Whitney<br>U test<br>p = 0.409 | Mann-Whitney<br>U test<br>p = 0.391 | Mann-Whitney<br>U test<br>p = 0.797 | Mann-Whitney<br>U test<br>p = 0.156 | Student T test<br>p = 0.074 |
| PN deep | 56 (n = 35 of 63) | $15 \pm 2.0$ | $3.5 \pm 0.2$ | $16 \pm 1.0$ | $3.5 \pm 0.4$ | $0.39 \pm 0.03$ |
| PN superficial | 56 (n = 53 of 94) | $20 \pm 3.0$ | $3.1 \pm 0.2$ | $15 \pm 0.9$ | $2.7 \pm 0.3$ | $0.51 \pm 0.04$ |
| Statistics | $\chi^2$ test<br>p = 0.946 | Mann-Whitney<br>U test<br>p = 0.306 | Mann-Whitney<br>U test<br>p = 0.051 | Mann-Whitney<br>U test<br>p = 0.314 | Mann-Whitney<br>U test<br>p = 0.083 | Mann-Whitney<br>U test<br>p = 0.072 |
|  | IPSC |  |  |  |  |  |
| cell type | connectivity (%) | amplitude (pA) | rise time (ms) | decay time (ms) | latency (ms) | success rate |
| CA2 PN | 35 (n = 19 of 55) | $197 \pm 41.3$ | $3.8 \pm 0.4$ | $25 \pm 1.2$ | $6.3 \pm 0.7$ | $0.55 \pm 0.06$ |
| CA3 PN | 57 (n = 16 of 28) | $145 \pm 23.4$ | $4.5 \pm 0.4$ | $25 \pm 1.2$ | $7.5 \pm 0.9$ | $0.54 \pm 0.05$ |
| Statistics | $\chi^2$ test<br>p = 0.134 | Mann-Whitney<br>U test | Student T test | Mann-Whitney<br>U test | Mann-Whitney<br>U test | Student T test |
|  |  | p = 0.870 | p = 0.203 | p = 0.896 | p = 0.303 | p = 0.893 |
| PN deep | 47 (n = 16 of 34) | 199 ± 40.6 | 3.8 ± 0.4 | 25 ± 1.4 | 7.2 ± 0.8 | 0.52 ± 0.07 |
| PN superficial | 47 (n = 26 of 55) | 167 ± 27.5 | 4.9 ± 0.4 | 26 ± 1.2 | 6.8 ± 0.7 | 0.50 ± 0.05 |
| Statistics | $\chi^2$ test<br>p = 0.987 | Mann-Whitney<br>U test<br>p = 0.258 | Student<br>T test<br>p = 0.047* | Student<br>T test<br>p = 0.564 | Student<br>T test<br>p = 0.706 | Student<br>T test<br>p = 0.796 |

### PNs in area CA2 receive mixed excitatory and inhibitory responses from SuM input

Photostimulation of SuM input elicited excitatory post-synaptic potentials (EPSPs) of fairly small amplitude in area CA2 PNs held at −70 mV (Figure 1E and 1F). Because current clamp experiments also show that SuM stimulation input also recruits feedforward inhibition in area CA2 (Figure 1 C4 and D4), we asked if the amplitude of SuM input stimulation-evoked EPSPs in PNs could be controlled by inhibition. Interestingly, blocking inhibitory transmission with the GABA_A_ and GABA_B_ receptor antagonists SR95531 and CGP55845A led to a significant increase of light-evoked EPSP amplitude recorded in area CA2 PNs (Figure 1F and 1G; amplitudes of the first response were 0.18 ± 0.05 mV in control and 0.24 ± 0.05 mV in SR95531 & CGP55845A, n = 14; Wilcoxon signed-rank tests, p = 0.004 for the first PSP, p = 0.013 for the second PSP, p < 0.001 for the third PSP). Thus, this result demonstrates a negative control of SuM-driven excitation by feedforward inhibition.

### Basket cells are strongly recruited by SuM inputs

Because the hippocampus hosts a variety of interneurons (INs) that are involved in controlling specific aspects of PN excitability, we wished to establish which kind of IN was targeted by the SuM input to area CA2. We performed whole-cell recordings from INs in this area and assessed post-synaptic excitatory responses to SuM axons stimulation in these cells (Figure 2). In contrast with previous reports of an exclusive innervation of PNs by SuM (Maglóczky et al., 1994), we observed robust light-evoked excitatory transmission from SuM axons in 35 out of 62 interneurons (INs) with soma located in SP. Following anatomical biocytin-streptavidin staining and reconstructions of recorded INs (allowing unequivocal identification in 48 neurons), we were able to classify INs based on their physiological properties, somatic location and axonal arborization location. We classified 22 cells as basket cells (BCs) because their axonal arborizations were restricted to SP (Figure 2A). BCs fired APs at high frequency either in bursts or continuously upon depolarizing current injection and showed substantial repolarizing sag current when hyperpolarized (Table 3). Light-evoked EPSCs and EPSPs were readily observed in the vast majority of BCs (Figure 2A and C, Table 4) and reached large amplitudes in some instances. An additional 26 INs with soma in SP were classified as non-BCs because their axon did not target SP (Figure 2B). In our recordings, these cells fired in bursts and showed little sag during hyperpolarizing current injection steps (Table 3). We consistently observed no or very minor light-evoked excitatory transmission onto non-BCs (Figure 2C, Table 4). Furthermore, we recorded from 17 INs that had soma in stratum oriens (SO) and 9 in stratum radiatum (SR). Like non-BCs, these INs did not receive strong excitation from SuM fibers (Table 4). This data is consistent with the conclusion that SuM input preferentially forms excitatory synapses onto basket cells in area CA2.

**Figure 2.**
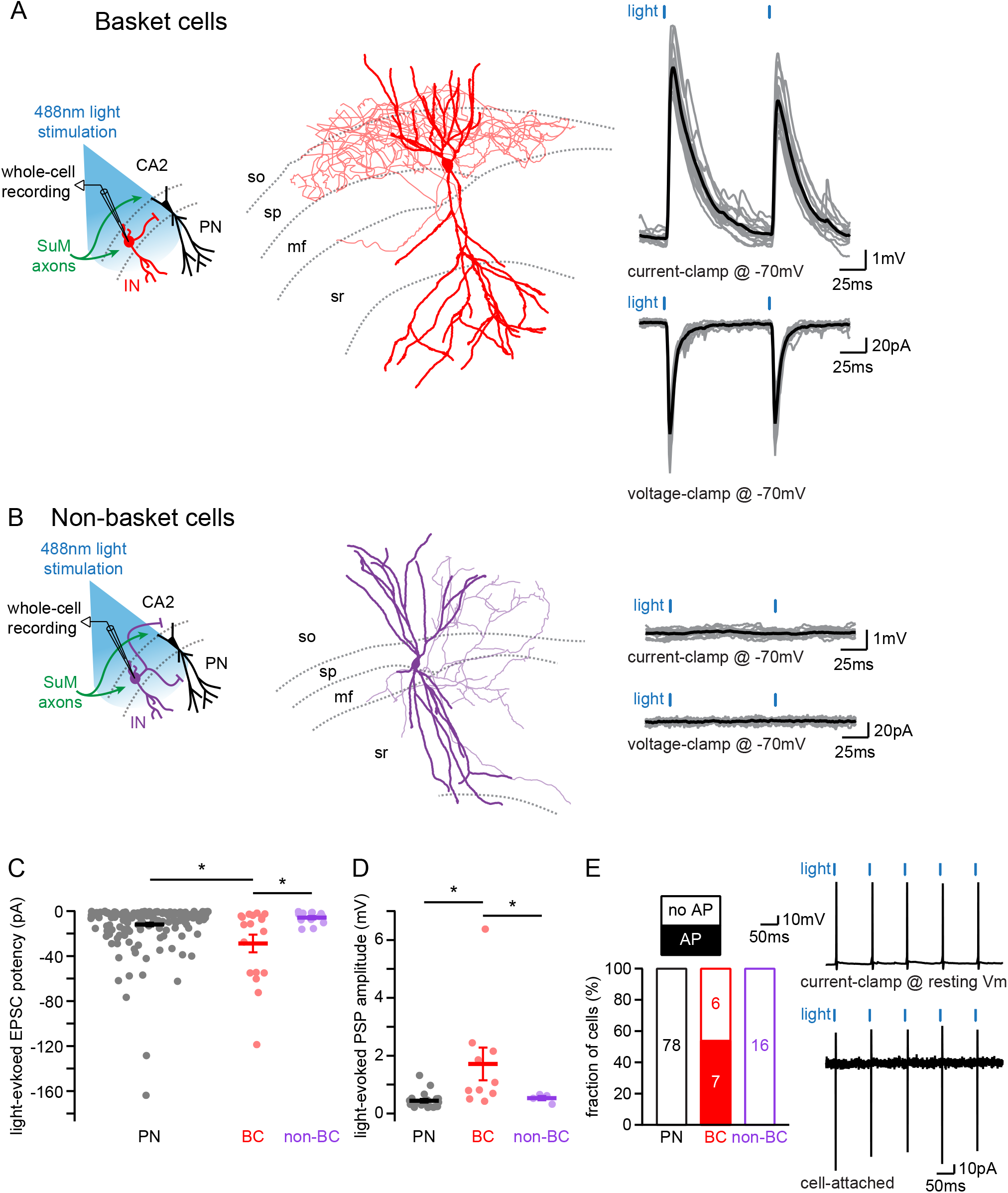
SuM input provides excitatory glutamatergic transmission to diverse population of PNs in area CA2. A-B. Left, diagrams illustrating whole-cell recordings in area CA2 and SuM fiber stimulation in acute slice preparation. Middle, example reconstruction of different cell types (soma and dendrites in thick lines, axon in thin lines, hippocampal strata in dotted grey lines). Right, sample traces of light-evoked EPSPs (top, individual traces in grey, average trace in black) and EPSCs (bottom, individual traces in grey, average trace in black). A. A Basket cell in area CA2. B. Non-basket cell. C. Summary graph of light-evoked EPSC potencies in PNs, BCs and non-BCs in area CA2 (individual cells shown as dots, population average shown as thick line, error bars represent SEM, PNs : n = 166; BC INs: n = 18; non-BCs: n = 13; Kruskal-Wallis test with Dunn-Holland-Wolfe post hoc test, p = 0.022). D. Summary graph of light-evoked PSP amplitudes in PNs, BCs and non-BCs (individual cells shown as dots, population average shown as thick line, error bars represent SEM, PNs : n = 20; BCs : n = 10; non-BCs : n = 4; Kruskal-Wallis test with Dunn-Holland-Wolfe post hoc test, p < 0.001). E. Left, proportion of post-synaptic CA2 PNs, BCs and non-BCs firing action potentials time-locked to light stimulation of SuM input. Right, sample traces of light-evoked action potentials in a BC recorded in current-clamp at resting membrane potential (top) and in cell-attached (bottom) configurations.

**Table 3.** Electrophysiological properties of interneurons in SuM-innervated area.

| | $V_M$ (mV) | $R_M$ (MOhm) | $C_M$ (pF) | firing adaptation index | sag (mV) |
| --- | --- | --- | --- | --- | --- |
| Basket cell (n = 16) | $-57.3 \pm 1.38$ | $144 \pm 28.1$ | $64.0 \pm 8.70$ | $0.74 \pm 0.05$ | $9.4 \pm 1.0$ |
| non-Basket Cell (n = 12) | $-55.6 \pm 1.84$ | $224 \pm 46.8$ | $52.0 \pm 5.90$ | $0.57 \pm 0.06$ | $5.9 \pm 1.4$ |
| interneuron SO (n = 6) | $-57.0 \pm 3.16$ | $201 \pm 21.0$ | $44.7 \pm 5.31$ | $0.61 \pm 0.11$ | $7.6 \pm 1.9$ |
| interneuron SR (n = 8) | $-60.1 \pm 2.89$ | $282 \pm 49.8$ | $39.6 \pm 3.18$ | $0.65 \pm 0.09$ | $8.1 \pm 2.1$ |
| Statistics | 1-way ANOVA test<br>p = 0.527 | 1-way ANOVA test<br>p = 0.100 | Kruskal-Wallis test<br>p = 0.354 | 1-way ANOVA test<br>p = 0.238 | 1-way ANOVA test<br>p = 0.292 |

**Table 4.** Characteristics of excitatory SuM light-evoked transmission onto interneurons & pyramidal cells.

| cell type | connectivity (%) | amplitude (pA) | rise time (ms) | decay time (ms) | latency (ms) | success rate |
| --- | --- | --- | --- | --- | --- | --- |
| Pyramidal Cell | 63 (n = 166 of 263) | $19 \pm 1.6^*$ | $3.4 \pm 0.1^*$ | $15 \pm 0.5^*$ | $2.9 \pm 0.1$ | $0.46 \pm 0.02$ |
| Basket Cell | 82 (n = 18 of 22) | $43 \pm 8.7^*$ | $1.7 \pm 0.3^*$ | $8.4 \pm 1.3^*$ | $3.1 \pm 0.4$ | $0.59 \pm 0.07$ |
| non-Basket Cell<br>interneuron SO<br>interneuron SR | 39 (n = 10 of 26)<br>12 (n = 2 of 17)<br>11 (n = 1 of 9) | $16 \pm 2.8$ | $2.6 \pm 0.5$ | $12 \pm 1.4$ | $3.4 \pm 0.7$ | $0.36 \pm 0.06$ |
| Statistics | $\chi^2$ test<br>$p = 0.006^*$ | Kruskal-Wallis test<br>$p = 0.016$<br>Dunn-Holland-Wolfe post hoc<br>$p < 0.05^*$ | 1-way ANOVA test<br>$p < 0.001$<br>Tukey post hoc<br>$p < 0.001^*$ | 1-way ANOVA test<br>$p < 0.001$<br>Tukey post hoc<br>$p < 0.001^*$ | 1-way ANOVA test<br>$p = 0.580$ | 1-way ANOVA test<br>$p = 0.066$ |

To fully assess the strength of SuM inputs onto the different cell types, we examined the following parameters for each population: the connectivity, success rate, amplitude, potency, kinetics, and latencies of EPSCs as well as the resulting depolarization of the membrane potential. First, SuM inputs preferentially innervated BCs as evidenced by a higher connectivity of EPSCs in BCs than in PNs or other INs (Table 4). Importantly, excitatory responses had short latencies with limited jitter (Table 4) indicating that the connection was monosynaptic in all cell types. When voltage-clamping cells at −70 mV, light-evoked EPSCs could be compared between different cell populations. However, not every photostimulation gave rise to an EPSC leading to an average success rate that tended to be highest in BCs (Table 4). In addition, BCs appeared to receive more excitation from SuM inputs than other cells types, as the amplitude of EPSCs were larger in BCs than in PNs (Table 4). EPSCs recorded in BCs also had faster kinetics than in PNs (Table 4). Interestingly, combining the success rate of EPSCs with their respective amplitudes to compute the potency of the SuM synapses revealed that it was significantly larger in BCs than in PNs and non-BCs (Figure 2C; potencies were 12 ± 1.6 pA for PNs, n = 166; 29 ± 7.8 pA for BCs, n = 18; 5.9 ± 1.5 pA for non-BCs, n = 13; Kruskal-Wallis test with Dunn-Holland-Wolfe post hoc test, p = 0.022). Consequently, EPSPs recorded at −70 mV were of larger amplitude in BCs than in PNs and non-BCs (Figure 2D; amplitudes were 0.44 ± 0.06 mV for PNs, n = 20; 1.71 ± 0.57 mV for BCs, n = 10; 0.53 ± 0.07 mV for non-BCs, n = 4; Kruskal-Wallis test with Dunn-Holland-Wolfe post hoc test, p < 0.001). When recording cell-attached or current-clamping BCs at their resting membrane potential (V_M_), photostimulation of SuM axons was able to evoke AP firing (Figure 2E) in multiple instances (n = 7 of 13), this was never observed in PNs (n = 0 of 78), non-BCs (n = 0 of 16), SR INs (n = 0 of 9) or SO INs (n = 0 of 8). These results show that SuM projections to area CA2 preferentially provide excitation to BCs that are likely responsible of the feedforward inhibition observed in PNs. This is in accordance with an efficient control of area CA2 PNs excitation by the SuM inhibitory drive as axons from BCs deliver the feedforward inhibition to the peri-somatic region of PNs.

### Parvalbumin-expressing basket cells mediate the feedforward inhibition recruited by SuM

In the hippocampus, BCs express either cholecystokinin (CCK) or parvalbumin (PV) (Klausberger and Somogyi, 2008). We found that in response to a 1 second depolarizing pulse, most BCs that received strong SuM excitatory input displayed very fast AP firing with little accommodation in the AP firing frequency (Table 3, Figure 3A and B). This firing behavior is similar to what has been reported for fast spiking PV-expressing BCs in CA1 (Pawelzik et al., 2002). In contrast, CCK-expressing BCs show a lower firing frequency and more accommodation during the train (Pawelzik et al., 2002). This result suggests that BCs connected by the SuM may be expressing PV. To directly confirm this hypothesis, we performed post-hoc immunostaining of recorded interneurons that received strong excitation from SuM input. Because of the dialysis inherent to the whole-cell recording conditions, we encountered difficulty staining for multiple cells. However, PV-immunoreactivity could unequivocally be detected in either the soma or dendrites of 7 connected BCs (Figure 3C). Therefore, this data demonstrates that at least a fraction of the recorded BCs connected by the SuM are expressing PV.

**Figure 3.**
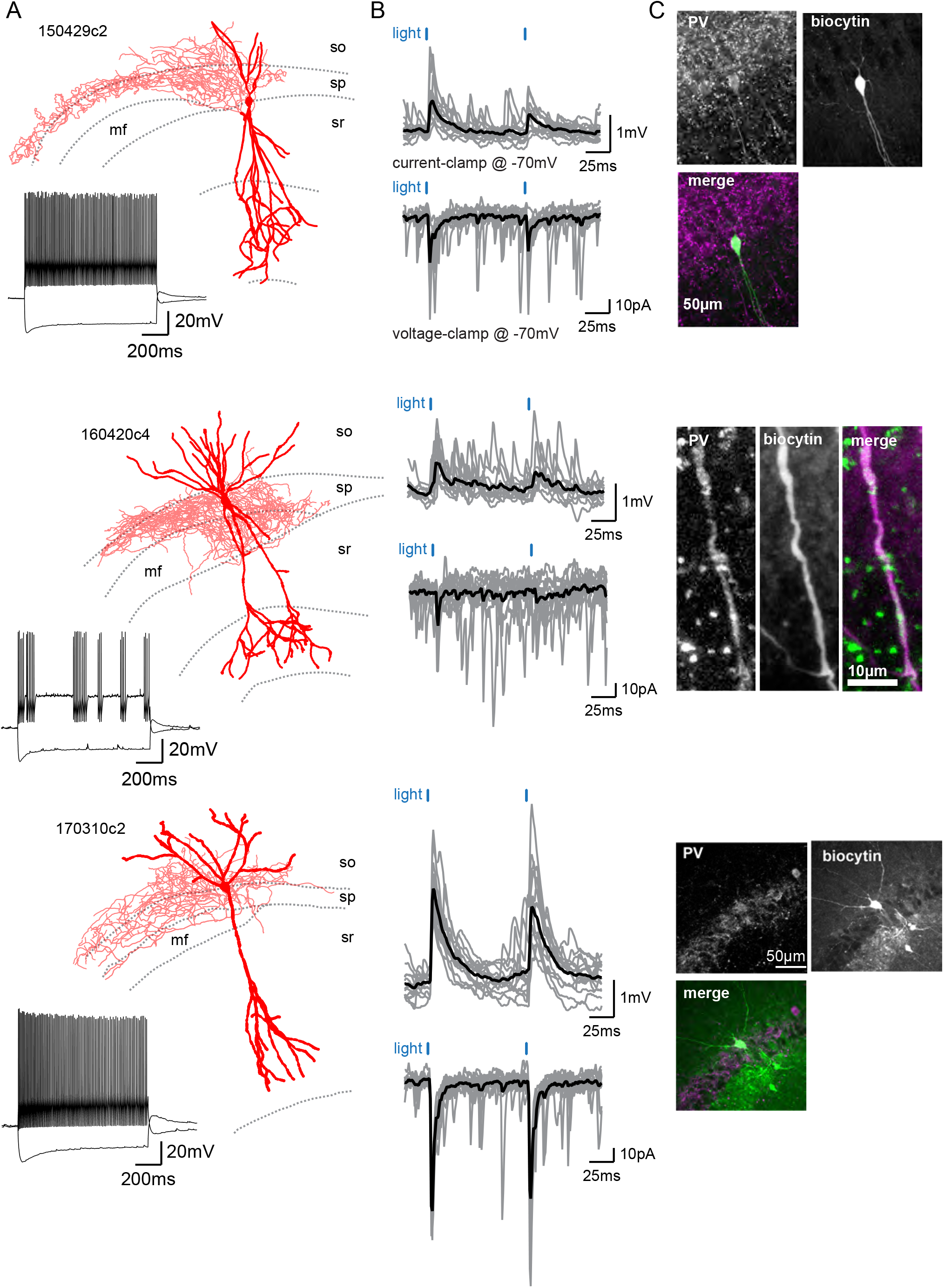
SuM inputs provide excitation to Parvalbumin-expressing BCs. A. Three biocytin reconstructions of BC INs with dendrites in red and axons in light red. Inset, current clamp steps to −400 pA and +400 pA display high-frequency AP firing and repolarizing sag current. B. Corresponding light-evoked EPSCs and EPSPs for the three reconstructed neurons (individual traces in grey, average trace in black). C. Corresponding PV immunostaining of the three interneurons: parvalbumin staining, biocytin labeling of the recorded cell, and merge (PV in magenta and biocytin in green).

Hence, to address whether the lack of PV staining in some cells was a consequence of dialysis or resulted from the fact that non-PV BC are also connected, we made use of a different strategy to differentiate PV and CCK INs. It has previously been demonstrated that PV+ BC transmission can be strongly attenuated by mu opioid receptor activation (MOR) while CCK+ BC transmission is insensitive to MOR activation (Glickfeld et al., 2008). Thus, in order to determine if SuM inputs preferentially target one subpopulation of BCs, we recorded from PNs in area CA2 and examined the sensitivity of light-evoked IPSCs to the application of the MOR agonist DAMGO (Figure 4A). We found that there was a near complete block of the light-evoked IPSC amplitude following 1 μM DAMGO application (Figure 4A; IPSC amplitudes were 343 ± 123 pA in control and 31 ± 12.4 pA in DAMGO hence a 88 ± 5.0 % block by DAMGO, n = 6 PNs; Wilcoxon signed-rank test, p = 0.031), while direct excitatory transmission remained unaffected (Figure 4A; EPSC amplitudes were 6.7 ± 1.1 pA in SR95531 & CGP55845A and 5.6 ± 0.9 pA after DAMGO, n = 17 PNs; Wilcoxon signed-rank test, p = 0.19).

**Figure 4.**
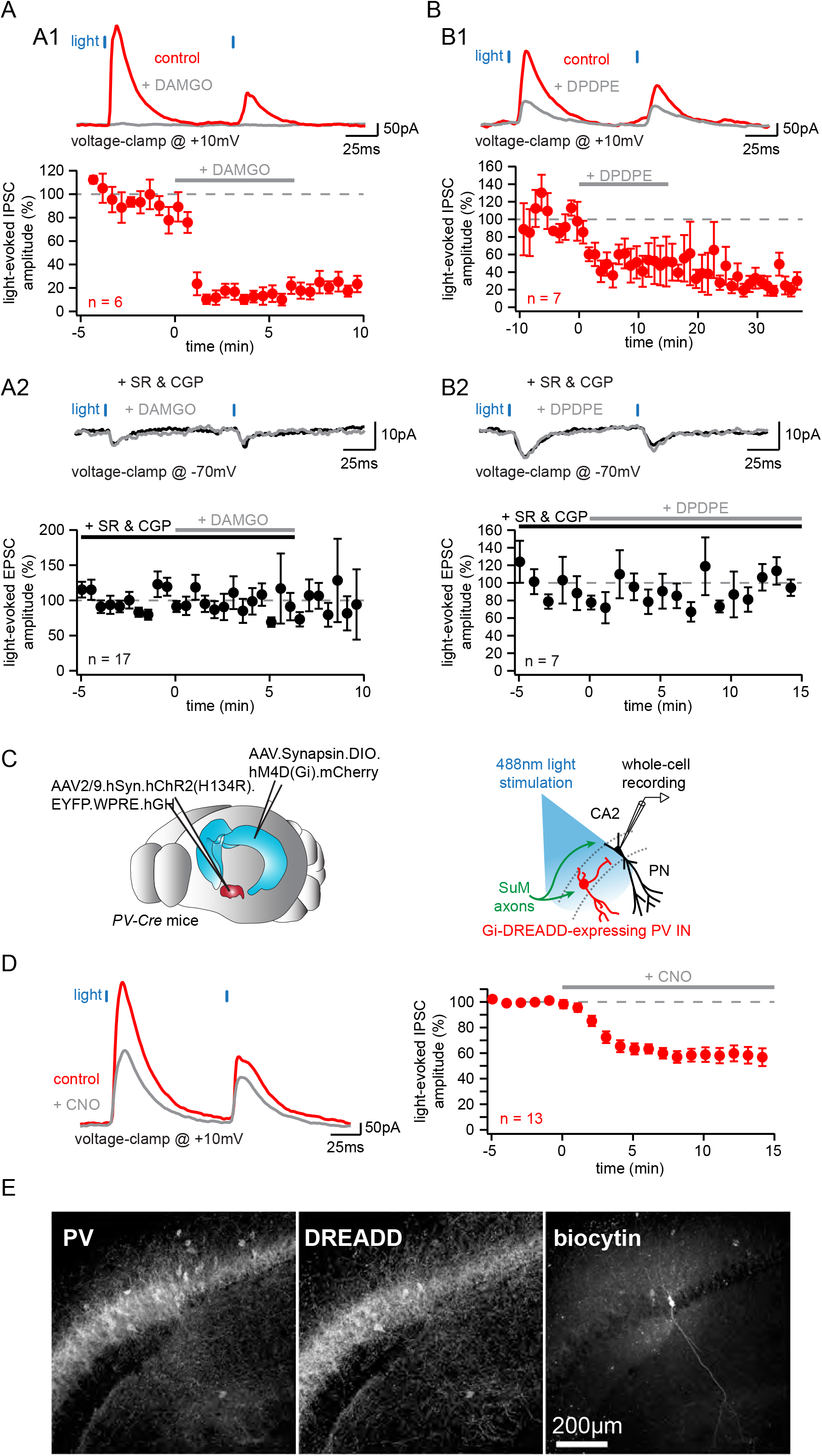
Parvalbumin-expressing BCs mediate the feedforward inhibition recruited by photostimulation of SuM fibers. A. Application of the mu-opioid receptor agonist, DAMGO, results in the complete abolition of light-evoked SuM inhibitory transmission. A1, sample traces (top, control in red, DAMGO in grey) and summary graph of light-evoked IPSC amplitudes recorded in area CA2 PNs before and after application of 1 μM DAMGO (bottom, n = 6, error bars represent SEM). A2, sample traces (top, SR95531 & CGP55845A in black, DAMGO in grey) and summary graph of light-evoked EPSC amplitudes recorded in area CA2 PNs before and after application of 1μM DAMGO (bottom, n = 17, error bars represent SEM). B. Application of the delta-opioid receptor agonist, DPDPE, results in the long-term depression of light-evoked SuM inhibitory transmission. B1, sample traces (top, control in red, DPDPE in grey) and summary graph of light-evoked IPSC amplitudes recorded in area CA2 PNs before and after application of 0.5 μM DPDPE (bottom, n = 7, error bars represent SEM). B2, sample traces (top, SR95531 & CGP55845A in black, DAMGO in grey) and summary graph of light-evoked EPSC amplitudes recorded in area CA2 PNs before and after application of 0.5 μM DPDPE (bottom, n = 7, error bars represent SEM). C. Left, diagrams illustrating the method to infect SuM neurons and selectively inhibit PV+ INs in area CA2. An AAV allowing the Cre-dependent expression of inhibitory DREADD was injected bilaterally into area CA2 of the dorsal hippocampus and another AAV allowing the expression of ChR2 was injected into the SuM of PV-Cre mice, allowing optogenetic stimulation of SuM inputs and pharmacogenetic inhibition of PV+ INs by application of the DREADD agonist CNO at 10 μM. Right, diagram of the recording configuration. D. Silencing of PV+ INs by inhibitory DREADDs reduces SuM feedforward inhibition onto area CA2 PNs. Sample traces (left, control in red, CNO in grey) and summary graph (right) of light-evoked IPSC amplitudes recorded in CA2 PNs before and after application of 10μM CNO (n = 13, error bars represent SEM). E. Example immunostaining against PV and DREADD with biocytin labelling in area CA2 from a slice used in these experiments.

Because a fraction of PV+ INs in area CA2 is also the substrate of an iLTD of feedforward inhibition from CA3 mediated by delta opioid receptor (DOR) activation, we sought to further refine our characterization of the SuM feedforward inhibition by assessing its sensitivity to DOR activation. Application of 0.5 µM of the DOR agonist DPDPE led to a long-term reduction of light-evoked IPSCs recorded in area CA2 PNs, similar to the iLTD seen by CA3 input stimulation (Figure 4B; amplitudes were 168 ± 28 pA in control and 64 ± 22 pA in DPDPE hence a 61 ± 14 % block by DPDPE, n = 7; paired-T test, p = 0.015), while leaving direct EPSCs unaffected (Figure 4B; amplitudes were 4.0 ± 1.6 pA in SR95531 & CGP55845A and 3.1 ± 1.1 pA after DPDPE, n = 7; Wilcoxon signed-rank test, p = 0.22). Further confirming the PV+ nature of INs responsible for the SuM feedforward inhibition, this result reveals that both the local CA3 and long-range SuM inputs converge onto the same population of INs to inhibit area CA2 PNs, thus enabling cross-talk between these routes through synaptic plasticity of PV+ INs.

Following up on this observation, we wished to genetically confirm that PV+ INs are responsible for the SuM feedforward inhibition over area CA2 PNs. As the dichotomy between PV+ versus CCK+ INs sensitivity to opioids has not been directly verified in area CA2, we used inhibitory DREADD to selectively inhibit PV+ INs in area CA2 while monitoring feedforward IPSCs from area CA2 PNs in response to SuM stimulation. To achieve that, we injected AAVs expressing a Cre-dependent h4MDi inhibitory DREADD in area CA2 of PV-Cre mice together with AAVs expressing ChR2 with a pan-neuronal promoter in the SuM (Figure 4C). We observed a substantial reduction of SuM-evoked IPSC amplitude recorded in area CA2 PNs upon application of 10 µM of the DREADD ligand CNO (Figure 4D; amplitudes were 847 ± 122 pA in control and 498 ± 87 pA in CNO hence a 42 ± 6.0 % block by CNO, n = 13; paired-T test, p < 0.001). Although we never measured a complete block of inhibitory responses, this result unequivocally places PV+ INs as mediators of the SuM feedfoward inhibition of area CA2 PNs. The incomplete block of IPSCs in these experiments could be a consequence of partial infection of PV+ INs in area CA2 by AAVs carrying DREADDs (Figure 4E; fraction of PV+ INs expressing DREADDs in CA2 = 75 ± 3.5 %, n = 13) and partial silencing of DREADD-expressing PV+ INs by CNO. Altogether, these combined results strongly indicate that SuM axons are efficiently and selectively exciting PV+ BCs in area CA2, thus driving a feedforward inhibition onto neighboring PNs.

### The feedforward inhibitory drive from SuM controls pyramidal neurons excitability

Given SuM axonal stimulation triggers an excitatory-inhibitory sequence in post-synaptic PNs, we asked which effect would prevail on PN excitability. In order to assess this, we mimicked an active state in PNs by injecting constant depolarizing current steps sufficient to sustain AP firing during 1 second while photostimulating SuM axons at 10 Hz (Figure 5A and 5B). We observed that recruitment of SuM inputs significantly delayed the onset of the first AP (Figure 5C; latency to the first AP were 221 ± 19.9 ms in control and 233 ± 19.1 ms with photostimulation, hence a 12.1 ± 4.3 ms increase upon photostimulation, n = 12; paired-T test, p = 0.016). In addition, given SuM neurons display theta-locked firing *in vivo*, we asked if rhythmic inhibition driven by SuM inputs in area CA2 could pace AP firing in PNs by defining windows of excitability. Indeed, photostimulation of SuM axons at 10 Hz led to a significant decrease of variability in the timing of AP firing by PNs (Figure 5D and 5E; standard deviations of the first AP timing were 36.9 ± 11 ms in control and 24.7 ± 7.4 ms with photostimulation, hence a 12.3 ± 5.3 ms decrease upon photostimulation, n = 12; Wilcoxon signed-rank tests, p < 0.001 for the first AP, p = 0.008 for the second AP, p = 0.004 for the third AP). Both the delay of AP onset and the reduction of AP jitter stemmed from the feedforward inhibition recruited by SuM inputs as application of GABA_A_ and GABA_B_ receptor antagonists abolished these effects of SuM stimulation (Figure 5C-E; latency to the first AP were 232 ± 19.8 ms in SR95531 & CGP55845A and 235 ± 18.0 ms with photostimulation, n = 6; Wilcoxon signed-rank test, p = 0.44; standard deviations of the first AP timing were 11.9 ± 2.0 ms in SR95531 & CGP55845A and 7.1 ± 1.5 ms with photostimulation, n = 6; Wilcoxon signed-rank tests, p = 0.22 for the first AP, p = 0.16 for the second AP, p = 0.09 for the third AP). These results reveal that the purely glutamatergic SuM input, by recruiting feedforward inhibition, has an overall inhibitory effect on PN excitability and can influence the timing and jitter of area CA2 PN action potential firing.

**Figure 5.**
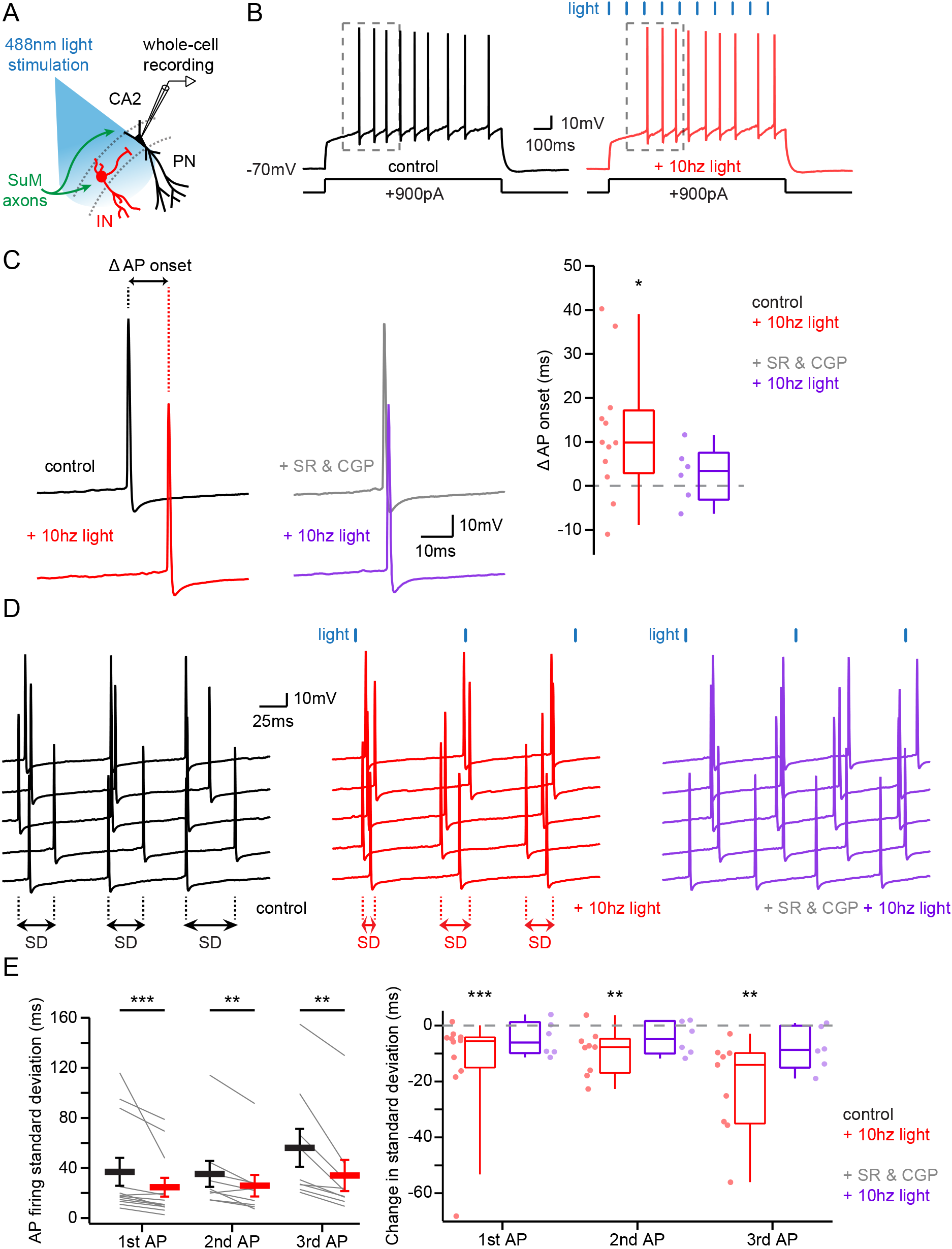
Area CA2 PNs receive a net inhibitory drive from SuM that controls AP firing properties. A. Diagram illustrating whole-cell recordings of area CA2 PNs and SuM fiber light stimulation in acute slice preparation. B. Example traces of a CA2 PN action potential firing in response to current injection in the absence (black traces) or presence of 10 Hz photostimulation of SuM inputs (red traces). C. Action potential onset is increased with 10 Hz SuM input photostimulation. Left, sample traces of the first AP in control and with inhibition blocked by 1 μM SR95531 & 2 μM CGP55845A application (light-off in black, light-on in red, light-off in SR95531 & CGP55845A in grey, light-on in SR95531 & CGP55845A in purple). Right, summary graph of photostimulation-induced delay of AP firing in area CA2 PNs before and after application of SR95531 & CGP55845A (control shown in red, n = 12, paired-T test, p = 0.016; SR95531 & CGP55845A shown in purple, n = 6; Wilcoxon signed-rank test, p = 0.44; individual cells shown with dots, boxplot represents median, quartiles, 10^th^ and 90^th^ percentiles). D. Sample traces of AP firing in repeated trials (light-off in black, light-on in red, light-on in SR95531 & CGP55845A in purple; during experiment photostimulation was interleaved with control, but are grouped here for demonstration purposes). E. AP jitter in CA2 PNs is reduced by activation of SuM inputs. Left, summary graph of the standard deviation of AP firing with or without 10 Hz photostimulation (n = 12; Wilcoxon signed-rank test, p < 0.001 for the first AP, p = 0.008 for the second AP, p = 0.004 for the third AP; individual cells shown with thin lines, population average shown as thick line, error bars represent SEM). Right, photostimulation-induced reduction of AP firing standard deviation in control and in SR95531 & CGP55845A (control, n = 12; Wilcoxon signed-rank tests, p < 0.001 for the first AP, p = 0.008 for the second AP, p = 0.004 for the third AP; SR95531 & CGP55845A, n = 6; Wilcoxon signed-rank tests, p = 0.22 for the first AP, p = 0.16 for the second AP, p = 0.09 for the third AP; individual cells shown with dots, boxplot represents median, quartiles, 10^th^ and 90^th^ percentiles).

It has been reported that the AP discharge of SuM neurons in vivo is phase-locked to the hippocampal theta rhythm (Kocsis and Vertes, 1994). Because theta rhythm is a brain state characterized by elevated levels of acetylcholine, we approximately mimicked these conditions in the hippocampal slice preparation by bath application of 10 µM of the cholinergic agonist carbachol (CCh). Under these conditions, CA2 PNs depolarize and spontaneously fire rhythmic bursts of APs, and the properties of these AP bursts are tightly controlled by excitatory and inhibitory synaptic transmission (Robert et al., 2020). Of note, we observed that CCh application depressed the SuM-CA2 excitatory and inhibitory drive and decreased short-term depression at these synapses (Supplemental Figure 3). Under these conditions, we asked how this spontaneous AP bursting activity would be affected by activation of the SuM input by triggering 10 second-long trains of 0.5 ms light pulses delivered at 10 Hz to stimulate SuM axons at the onset of bursts (Figure 6A). Because of the intrinsic cell-to-cell variability of bursting kinetics, we photo-stimulated SuM inputs only during interleaved bursts in the same cells. To do this, bursts were detected automatically with an online threshold detection system that started the photostimulation pulse train after the first AP of every alternating burst, starting with the second burst (Figure 6A and B). For analysis, the number of APs and bursting kinetics could be compared within the same cell. We observed a significant decrease in the number of APs fired during a burst when SuM inputs were photo-stimulated as compared to interleaved control bursts (Figure 6C and 6D; numbers of APs per burst were 15.2 ± 2.3 in control and 6.9 ± 1.3 with photostimulation, n = 7; paired-T test, p = 0.031). In control bursts, the AP firing rate of CA2 PNs initially increases, and then progressively decreases. In the photo-stimulation bursts, the initial increase of AP firing frequency was absent, and the subsequent AP firing frequency was reduced (Figure 6E; 2-way ANOVA on firing rate over time in light-on vs light-off conditions; light factor, p < 0.001; time factor, p < 0.001; light × time factor, p = 0.052).

**Figure 6.**
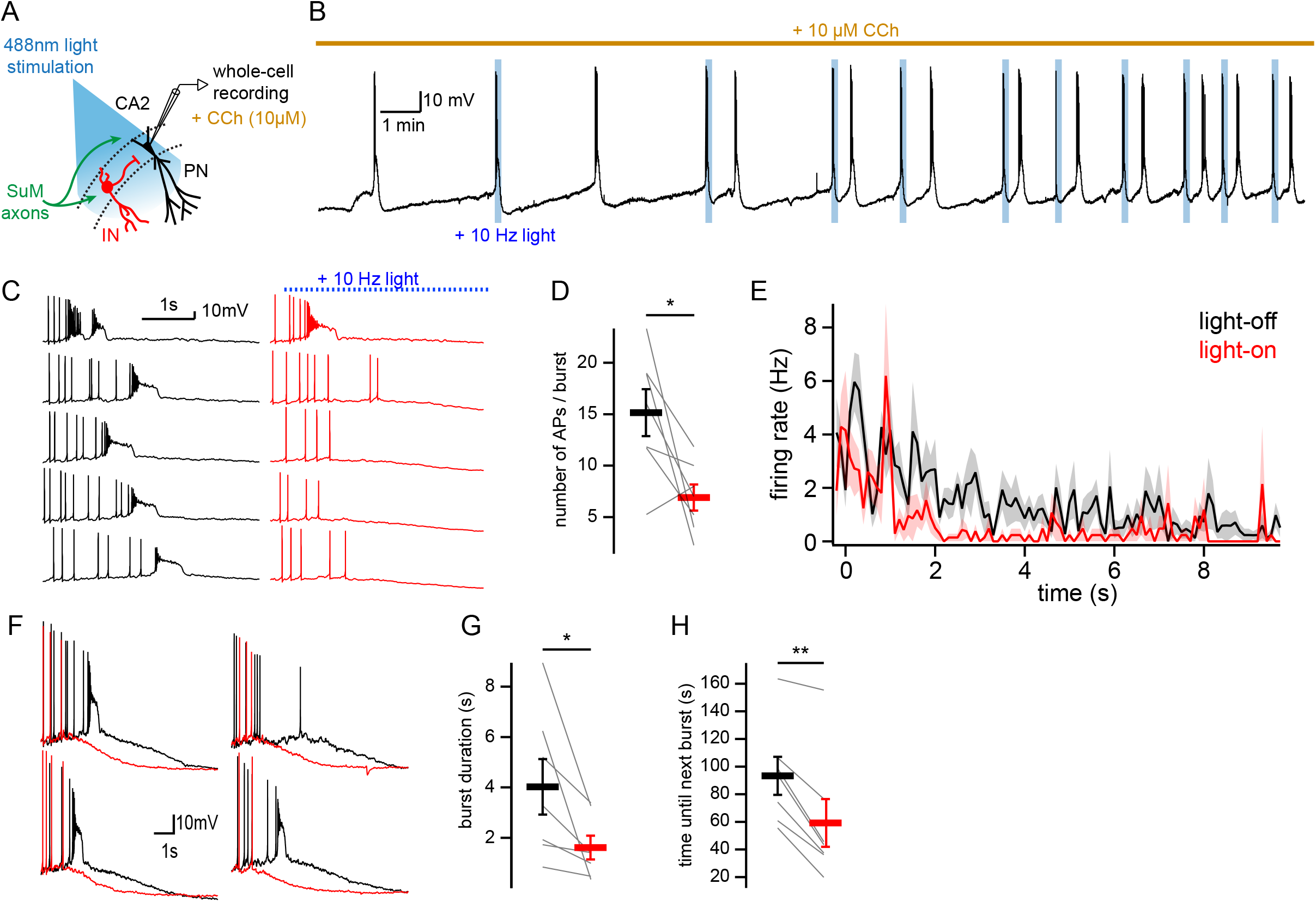
SuM input shapes CA2 PN AP bursts in conditions of elevated cholinergic tone. A. Diagram illustrating whole-cell recordings of area CA2 PNs with light stimulation of SuM fibers in an acute slice preparation. B. Sample trace of spontaneous AP bursting activity recorded from a CA2 PN during bath application of 10 μM CCh. For every even-numbered burst, a 10 Hz photostimulation (blue bars) was delivered to excite SuM inputs in area CA2 allowing a comparison of burst AP firing in the same cell. C. Sample traces of AP firing during bursts for light-off (left, black) and light-on (right, red) epochs. D. Comparison of AP number / burst for light-off (black) and light-on (red) events (n = 7; individual cells shown as thin lines, population average shown as thick line, error bars represent SEM; paired-T test, p = 0.031). E. Average firing rate during spontaneous burst events with SuM photostimulation (red, light-on) and controlled inter-leaved burst events (black, light-off). Shaded area represents SEM for 7 cells each with between 3 and 13 bursts analyzed in light-on and light-off conditions (2-way ANOVA, light factor: p < 0.001, time factor: p < 0.001, light x time factor: p = 0.052). F. Example burst events with (red) and without (black) SuM photostimulation overlayed and on a scale that shows the rapidly hyperpolarizing membrane potential that occurs with SuM input stimulation. G. Comparison of bursts duration for events with (red) and without (black) photostimulation (n = 7; individual cells shown as thin lines, population average shown as thick line, error bars represent SEM; paired-T test, p = 0.037). H. Comparison of time elapsed to next burst onset following bursts with (red) or without (black) photosimulation (n = 7; individual cells shown as thin lines, population average shown as thick line, error bars represent SEM; paired-T test, p = 0.001).

In the presence of CCh, spontaneous AP bursting is preceded by a membrane depolarization. Following several seconds of AP firing, the membrane potential of CA2 PNs remains depolarized for several seconds, and slowly hyperpolarizes until the next burst event. We observed that photo-stimulation of SuM inputs resulted in a striking reduction in the amount of time the membrane potential remained depolarized, and this is likely why the burst duration was significantly shorter in bursts with SuM photo-stimulation (Figure 6F and G; burst duration was 4.0 ± 1.1 s in control and 1.6 ± 0.5 s with photostimulation, n = 7; paired-T test, p = 0.037). The rate and level of V_M_ repolarization following bursts were not significantly changed by SuM input photostimulation (V_M_ repolarization rate was −3.3 ± 0.6 mV/s in control and −3.6 ± 0.7 mV/s with photostimulation, n = 7; paired-T test, p = 0.601; post-burst V_M_ was −62.8 ± 1.7 mV in control and −62.0 ± 2.0 mV with photostimulation, n = 7; paired-T test, p = 0.173), however the inter-burst time interval was reduced. Indeed, AP bursts with SuM input activation were followed more rapidly by another burst of AP than the ones without SuM input activation (Figure 6B, H; time until next burst was 93 ± 14 s in control and 59 ± 17 s with photostimulation, n = 7; paired-T test, p = 0.001), which could be due to both short-term depression of inhibitory transmission after repeated activation during the SuM input photostimulation train and reduced activation of hyperpolarizing conductances during bursts shortened by SuM input photostimulation. Thus, in our preparation, SuM input activation is able to modify the spontaneous bursting activity of CA2 PNs under conditions of high cholinergic tone.

As SuM input controls burst firing of action potentials and likely paces activity in area CA2, we wondered how the subsequent output of CA2 PNs would affect their post-synaptic targets. Because CA2 PNs strongly project to CA1 PNs, this activity is likely to influence CA1 encoding and hippocampal output. Thus, we examined the consequences of SuM-CA2 input stimulation on area CA1 both in vivo and in acute slices treated with CCh to induce spontaneous activity (Figure 7).

**Figure 7.**
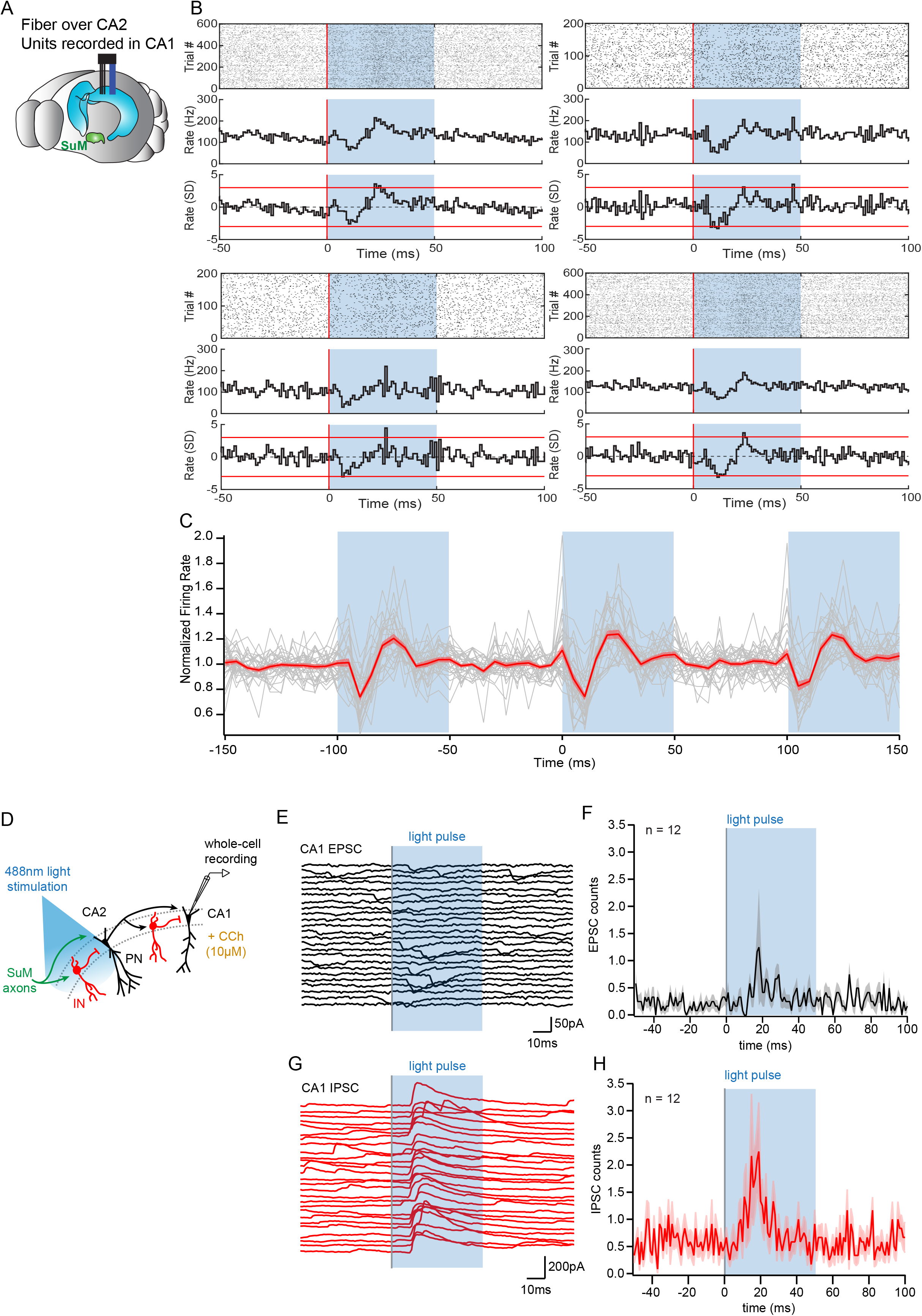
Consequences of SuM input on area CA2 output to CA1. A. Diagram illustrating in vivo recording in CA1 with tetrodes and SuM axon terminals stimulation over CA2 with an implanted optical fiber. B. Representative data from 4 multi-unit recordings. Raster plot (top) showing CA1 AP firing activity before and during photostimulation of SuM fibers in area CA2. The corresponding firing rate histogram (middle) of four tetrodes placed in the CA1 pyramidal cell layers, as well as plots of standard deviation (SD; bottom). Red lines indicate +/− 3SD. C. Individual (grey) and average (red) normalized firing rates from 31 multiunit recordings, 3 consecutive light stimulation epochs are displayed to help visualizing the consistency of the effect of SuM input light stimulation over area CA2 on CA1 multi-unit firing; the shaded area represents the SEM. D. Diagram illustrating whole-cell recordings of area CA1 PNs and SuM fiber light stimulation over area CA2 in acute slice preparation. E-H. Example waterfall plots (E, G) and corresponding peri-stimulus time histogram (F, H, population average shown as thick line, shaded area represents SEM) of EPSCs (black) and IPSCs (red) recorded from a CA1 PN ex vivo during photostimulation of SuM input over area CA2 with bath application of 10 μM CCh.

ChR2-EYFP was expressed in the SuM of Csf2rb2-cre mice in a cre-dependent manner and the mice were implanted with a microdrive targeting tetrodes to region CA1 and an optical fiber to the SuM terminals in CA2 (Figure 7A). Mice were placed in a small box (familiar context) and left free to explore as blue (473 nm) laser light pulses (50 ms pulse width) were applied to the SuM terminals at 10 Hz. Across 23 recording sessions in five mice we found that the activation of SuM terminals in CA2 resulted in a significant and reproducible change in the multiunit spiking activity recorded in the pyramidal cell layer of CA1 on 34 of 55 tetrodes. The firing rate change was similar across individual tetrodes (Figure 7B and C), with a decrease in the normalized firing rate starting shortly after laser onset and continuing for about 10 ms, followed immediately by a rebound-like increase to about 20 % greater than baseline firing rate (Figure 7B and C).

In order to get a better mechanistic understanding of this observation, we set out to decipher how SuM activity in area CA2 influences CA1 in the hippocampal slice preparation. To this end, we used the same photostimulation protocol used in vivo that consisted of light stimulation trains of 50 ms-long pulses delivered at 10 Hz for 1 second, repeated every 10 seconds for 2 minutes and interleaved with light-off sweeps of the same duration, with the microscope objective centered on area CA2. Whole-cell patch-clamp recordings of CA1 PNs were obtained in acute hippocampal slices superfused with CCh and subjected to this light stimulation protocol (Figure 7D). We asked what synaptic events may be responsible for the decreased firing of CA1 units observed 10 – 20 ms after light onset in vivo (Figure 7A-C). Whole-cell recordings of CA1 PNs showed an absence of EPSCs time-locked to the photostimulation in all but one case (n = 11/12) (Figure 7E and F). In contrast, we often (n = 7/12) observed light-evoked IPSCs in CA1 PNs occurring 10 – 20 ms after light onset (Figure 7G and H). Therefore, the reduction in firing of CA1 units in vivo is likely caused by increased inhibitory inputs onto CA1 PNs within 10 – 20 ms of SuM fiber stimulation over area CA2. This result highlights a contribution of SuM input to controlling CA2 output that regulate CA1 activity in vivo and provides a mechanistic interpretation of this observation at the circuit level.

## Discussion

In this study, we provide direct evidence for a functional connection between the hypothalamus and the hippocampus. Using stereotaxic injection of viral vectors in combination with transgenic mouse lines to express channelrhodopsin in a projection-specific manner, we have been able to selectively stimulate SuM axons in area CA2 of the hippocampus, allowing for the direct examination of synaptic transmission. This approach yielded novel functional physiological information about the SuM post-synaptic targets and overall consequences of activation. We found that, in contrast to previous anatomical reports, SuM inputs form synapses onto both PNs and INs in area CA2. The excitatory drive evoked by light-stimulation of SuM inputs was significantly larger for BC INs, which we demonstrate are likely PV+. The resulting feedforward inhibition recruited by SuM input stimulation enhanced the precision of AP timing of CA2 PNs in conditions of low and high cholinergic tone. The modified CA2 output evoked poly-synaptic inhibition in area CA1, likely responsible for a decrease firing rate of CA1 units in vivo. Overall, we demonstrate that SuM input controls CA2 output to area CA1 by recruiting feedforward inhibition.

### SuM inputs to area CA2 form a microcircuit where PV+ basket cells strongly inhibit pyramidal neurons

Glutamatergic innervation of area CA2 by the SuM has been previously described by tracing studies (Kiss et al., 2000; Soussi et al., 2010) and presumed to form synapses exclusively onto PNs (Maglóczky et al., 1994). Our experimental strategy allowed for the direct examination of the post-synaptic targets of SuM glutamatergic axons. Our results confirm that PNs in area CA2 indeed receive excitatory synapses from SuM axons. However, in contrast to what had been proposed in previous studies, we observed that SuM inputs target not only PNs but also INs in area CA2. Importantly, we identified a specific subpopulation of INs as PV+ BCs which were the cell type most potently excited by SuM. These BCs could fire action potentials upon SuM inputs photostimulation leading to a substantial feedforward inhibition of neighboring PNs. Consistent with the perisomatic targeting of BCs axons, recruitment of BCs by SuM resulted in the control of PNs excitability. This finding demonstrates that SuM activity can pace action potential firing in PNs through recruitment of PV+ BCs. The inhibitory action of the SuM input to area CA2 contrasts with the overall excitatory effect of the SuM-DG path (Hashimotodani et al., 2018; Li et al., 2020; Mizumori et al., 1989; Nakanishi et al., 2001).

### Consequences of SuM input on area CA2 output

Recent work has demonstrated a strong excitatory drive from area CA2 to CA1 (Chevaleyre and Siegelbaum, 2010; Kohara et al., 2014; Nasrallah et al., 2019). Consequently, modification of CA2 output through synaptic plasticity (Nasrallah et al., 2019) or neuromodulation (Tirko et al., 2018) affects CA1 activity. This observation is critical when considering social memory formation, which is known to depend on CA2 output (Hitti and Siegelbaum, 2014; Stevenson and Caldwell, 2014) and is likely encoded in downstream ventral CA1 (Okuyama et al., 2016). CA2-targeting cells in the SuM have recently been shown to be highly active during novel social exploration (Chen et al., 2020). From our results, we hypothesize that this novel social signal from the SuM, acts via the PV+ inhibitory network in area CA2 to control the timing of CA2 output onto area CA1.

The population of INs potently excited by SuM transmission display many features that allow us to classify them as PV+ BCs. They have somas located in the somatic layer, have densely packed perisomatic-targeted axons, are fast spiking, show PV immuno-reactivity, are sensitive to MOR and DOR activation, and their selective silencing reduces SuM driven feed-forward inhibition of area CA2 PNs. Recent studies have indicated that DOR-mediated inhibitory synaptic plasticity of PV+ INs in area CA2 is required for social recognition memory (Domínguez et al., 2019) and further, that exposure to a novel conspecific induces a DOR-mediated plasticity in this same inhibitory network in area CA2 (Leroy et al., 2017) Thus, our finding that SuM input acts via PV+ interneurons fits with previous results, and provides a link between social novelty information and local hippocampal inhibitory plasticity.

By recruiting feedforward inhibition, SuM activity paces and temporally constrains AP firing from CA2 PNs undergoing depolarization. More critically, in conditions of elevated cholinergic tone relevant to SuM activity in vivo, CA2 PNs depolarize and fire bursts of APs that can be shaped by SuM input both by controlling AP firing as well as membrane depolarization. While this result was obtained by triggering SuM input stimulation to the onset of burst firing by CA2 PNs, in vivo and acute slice experiments revealed a consistent influence of CA1 PN AP firing by SuM input to area CA2 regardless of the timing of SuM input stimulation relative to CA2 PN AP burst firing. These results demonstrate a powerful control of SuM input over CA2 output when PNs are spontaneously firing bursts of APs, a firing mode that is most efficient at influencing CA1 activity (Tirko et al., 2018). Optogenetic experiments have recently shown that CA2 PNs can drive a strong feedforward inhibition in area CA1 (Nasrallah et al., 2019). Although SuM input likely does not directly drive feedforward inhibition in area CA1 (Chen et al., 2020), the recruitment of feedforward inhibition in area CA2 by SuM input activation could curtail the time window of spontaneous firing in CA2 PNs and effectively lead to a synchronized drive of feedforward inhibition by area CA2 over area CA1. We postulate that the concerted IPSC that we detect in area CA1 with SuM fiber photostimulation in area CA2 corresponds to the large decrease in firing that is observed in CA1 multi-unit recordings in vivo. Thus, these data provide evidence for a long-range control of CA2 bursting activity and the consequences in downstream area CA1 in conditions of high cholinergic tone that accompanies theta oscillations in vivo during which SuM is active.

### Gating of area CA2 activity by PV+ INs and significance for pathologies

The density of PV+ INs in area CA2 is strikingly higher than in neighboring areas CA3 and CA1 (Botcher et al., 2014; Piskorowski and Chevaleyre, 2013). This population of INs has been shown to play a powerful role in controlling the activation of CA2 PNs by CA3 inputs (Nasrallah et al., 2015). We show in this study that long-range inputs from the SuM can strongly recruit PV+ BCs, which in turn inhibit PNs in this area. Hence, both intra-hippocampal inputs from CA3 and long-range inputs from the SuM converge onto PV+ INs to control CA2 PN excitability and output.

Postmortem studies have reported losses of PV+ INs in area CA2 in pathological contexts including bipolar disorder (Benes et al., 1998), Alzheimer’s disease (Brady and Mufson, 1997), and schizophrenia (Benes et al., 1998; Knable et al., 2004). Consistent with these reports, in a mouse model of the 22q11.2 deletion syndrome, we found a loss of PV staining and deficit of inhibitory transmission in area CA2 that were accompanied by impairments in social memory (Piskorowski et al., 2016). We postulate that the PV+ INs altered during pathological conditions may be the same population of PV+ BCs recruited by long-range SuM inputs. Indeed, the DOR-mediated plasticity onto PV+INs is altered in the 22q11.2 deletion syndrome, and we show here that the PV+ INs targeted by the SuM also express DOR. Thus, the loss of function of PV+ INs in area CA2 could disrupt proper long-range connection between the hippocampus and the hypothalamus and possibly contribute to some of the cognitive impairments observed in schizophrenia animal models. Further, pharmacological mouse models of schizophrenia have reported increased c-fos immunoreactivity in the SuM as well as memory impairments (Castañé et al., 2015). Although several alterations in these models of schizophrenia could lead to deficits of hippocampal-dependent behavior, abnormalities of the SuM projection onto area CA2 appear as a potential mechanism that warrants further investigation.

## Materials & Methods

All procedures involving animals were performed in accordance with institutional regulations (French Ministry of Research and Education protocol #12406-2016040417305913). Animal sample sizes were estimated using power tests with standard deviations and ANOVA values from pilot experiments. A 15 % failure rate was assumed to account for stereotaxic injection errors and slice preparation complications. Every effort was made to reduce animal suffering.

### Use of the Tg(Slc17ab-icre)10Ki mouse line

we used the Tg(Slc17ab-icre)10Ki mouse line that was previously generated (Borgius et al., 2010) and expresses the Cre recombinase under the control slc17a6 gene coding for the vesicular glutamate transporter isoform 2 (VGluT2).

### Use of the csf2rb2-Cre mouse line

We used the csf2rb2-Cre mouse line that was recently generated (Chen et al., 2020) and expresses the Cre recombinase under control of the csf2rb2 gene that shows selective expression in the SuM.

### Use of the Pvalbtm1(cre)Arbr/J mouse line

we used the Pvalbtm1(cre)Arbr/J mouse line that was previously generated (Hippenmeyer et al., 2005) and expresses the Cre recombinase under the control Pvalbm gene coding for parvalbumin (PV).

### Stereotaxic viral injection

Animals were anaesthetized with ketamine (100 mg/kg) and xylazine (7 mg/kg). The adeno-associated viruses AAV9.EF1a.DIO.hChR2(H134R).EYFP and AAV9.hSynapsin.EGFP.WPRE.bGH were used at 3×10^8^ vg, the AAV.Synapsin.DIO.hM4D(Gi).mCherry was used at 3.6×10^9^ vg and the AAV2/9.hSyn.hChR2(H134R).EYFP.WPRE.hGH was used at 3.7×10^13^ vg. The retrograde tracer CAV2-cre virus was used at 2.5×10^12^ vg. 500 nL of virus was unilaterally injected into the brain of 4 week-old male wild type C57BL6, Tg(Slc17ab-icre)10Ki (VGluT2-Cre), csf2rb2-cre (SuM-Cre) or Pvalbtm1(cre)Arbr/J (PV-Cre) mice at 100 nL/min and the injection cannula was left at the injection site for 10 min following infusion. In the case of AAV.Synapsin.DIO.hM4D(Gi)-mcherry injection in PV-Cre mice, bilateral injections were performed in dorsal CA2. The loci of the injection sites were as follows: anterior–posterior relative to bregma: −2.8 mm for SuM, −1.6 mm for CA2; medial-lateral relative to midline: 0 mm for SuM, 1.9 mm for CA2; dorsal-ventral relative to surface of the brain: 4.75 mm for SuM, 1.4 mm for CA2.

### Electrophysiological recordings

Transverse hippocampal slices were prepared at least 3 weeks after viral injection and whole-cell patch-clamp recordings were performed from PNs and INs across the hippocampal CA regions. In the case of PV-Cre mice injected with AAV.Synapsin.DIO.hM4D(Gi)-mcherry, slices were prepared 6 weeks after viral injection. Animals were deeply anaesthetized with ketamine (100 mg/kg) and xylazine (7 mg/kg), and perfused transcardially with a N-methyl-D-glucamin-based (NMDG) cutting solution containing the following (in mM): NMDG 93, KCl 2.5, NaH_2_PO_4_ 1.25, NaHCO_3_ 30, HEPES 20, glucose 25, thiourea 2, Na-ascorbate 5, Na-pyruvate 3, CaCl_2_ 0.5, MgCl_2_ 10. Brains were then rapidly removed, hippocampi were dissected out and placed upright into an agar mold and cut into 400 μm thick transverse slices (Leica VT1200S) in the same cutting solution at 4 °C. Slices were transferred to an immersed-type chamber and maintained in artificial cerebro-spinal fluid (ACSF) containing the following (in mM) : NaCl 125, KCl 2.5, NaH_2_PO_4_ 1.25, NaHCO_3_ 26, glucose 10, Na-pyruvate 2, CaCl_2_ 2, MgCl_2_ 1. Slices were incubated at 32°C for approximately 20 min then maintained at room temperature for at least 45 min prior to patch-clamp recordings performed with either potassium- or cesium-based intracellular solutions containing the following (in mM): K- or Cs-methyl sulfonate 135, KCl 5, EGTA-KOH 0.1, HEPES 10, NaCl 2, MgATP 5, Na_2_GTP 0.4, Na_2_-phosphocreatine 10 and biocytin (4 mg/mL).

ChR2 was excited by 488 nm light delivered by a LED attached to the epifluorescence port of the microscope. Light stimulations trains consisted of 2-10 pulses, 0.5 ms long, delivered at 10 Hz, repeated every 20 s for at least 20 sweeps. For the patch-clamp recordings in area CA1 with stimulation of SuM axons in area CA2, 50 ms long light stimulation pulses were delivered every 10 seconds. We used a light intensity of 25 mW/mm^2^ which was experimentally determined as the lowest irradiance allowing TTX-sensitive maximal responses in all cell types and conditions. Data were obtained using a Multiclamp 700B amplifier, sampled at 10 kHz and digitized using a Digidata. The pClamp10 software was used for data acquisition. Series resistance were < 20 MOhm and were not compensated in voltage-clamp, bridge balance was applied in current-clamp. An experimentally determined liquid junction potential of approximately 9 mV was not corrected for. Pharmacological agents were added to ACSF at the following concentrations (in μM): 10 NBQX and 50 D-APV to block AMPA and NMDA receptors, 1 SR95531 and 2 CGP55845A to block GABA_A_ and GABA_B_ receptors, 1 DAMGO to activate δ-opioid receptors (MOR), 0.5 DPDPE to activate δ-opioid receptors (DOR), 10 clozapine N-oxide (CNO) to activate hM4D(Gi) DREADDs, 10 CCh to activate cholinergic receptors, 0.2 tetrodotoxin (TTX) to prevent sodic action potential generation.

### Surgery for *in vivo* recordings

All surgeries were performed in a stereotaxic frame (Narishige). Csf2rb2-cre male mice from 3 to 6 months of age were anaesthetized using 500□mg/kg Avertin. pAAV.DIO.hChR2(H134R).EYFP was injected into the SuM (−2.7 mm AP, +0.4 mm ML, −5.0 mm DV) using a 10□μL Hamilton microsyringe (701LT, Hamilton) with a beveled 33 gauge needle (NF33BL, World Precision Instruments (WPI)). A microsyringe pump (UMP3, WPI) with controller (Micro4, WPI) were used to set the speed of the injection (100 nl/min). The needle was slowly lowered to the target site and remained in place for 5□min prior to start of the injection and the needle was removed 10 min after infusion was complete. Following virus injection, a custom-built screw-driven microdrive containing six independently adjustable nichrome tetrodes (14 μm diameter), gold-plated to an impedance of 200 to 250 kΩ was implanted, with a subset of tetrodes targeting CA1, and an optic fiber (200 μm core diameter, NA=0.22) targeting CA2 (−1.9 mm AP, +/− 2.2 mm ML, −1.6 mm DV). Following recovery, the tetrodes were slowly lowered over several days to CA1 pyramidal cell layer, identified by characteristic local field potential patterns (theta and sharp-wave ripples) and high amplitude multiunit activity. During the adjustment period the animal was habituated every day to a small box in which recording and stimulation were performed.

### *In vivo* recording protocol

Recording was commenced following tetrodes reaching CA1. To examine the impact of SuM terminal stimulation in CA2 the mice were returned to the small familiar box and trains of 10 light pulses (473 nm, 10 mW/mm^2^ and pulse width 50 ms) were delivered to the CA2 at 10 Hz. The pulse train was repeated every 10 seconds for at least 20 times as the animals freely explored the box. Multiunit activity was recorded using a DigitalLynx 4SX recording system running Cheetah v.5.6.0 acquisition software (Neuralynx). Broadband signals from each tetrode were filtered between 600 and 6,000 Hz and recorded continuously at 32 kHz. Recording sites were later verified histologically with electrolytic lesions as described above and the position of the optic fiber was also verified from the track.

### *In Vivo* data analysis

Spike and event timestamps corresponding to onset of each laser pulse were imported into Matlab (MathWorks) and spikes which occurred 50 ms before and 100 ms after each laser pulse were extracted. Raster plots were generated using a 1 ms bin size. Similar results were obtained using 5 ms and 10 ms bin size (data not shown). Firing rate histograms were calculated by dividing total number of spikes in each time bin by that bin’s duration. Each firing rate histogram was normalized by converting it into z-score values. Mean standard deviation values for the z-score calculation were taken from pre-laser pulse time period. To average the response across all mice, for each tetrode the firing rate in each bin was normalized to the average rate in the pre-laser period.

### Immunochemistry and cell identification

Midbrains containing the injection site were examined post-hoc to ensure that infection was restricted to the SuM.

Post-hoc reconstruction of neuronal morphology and SuM axonal projections were performed on slices and midbrain tissue following overnight incubation in 4 % paraformaldehyde in phosphate buffered saline (PBS). Midbrain sections were re-sliced sagittally to 100 μm thick sections. Slices were permeabilized with 0.2 % triton in PBS and blocked overnight with 3 % goat serum in PBS with 0.2 % triton. Primary antibody (life technologies) incubation was carried out in 3 % goat serum in PBS overnight at 4°C. Channelrhodopsin-2 was detected by chicken primary antibody to GFP (Life technologies) (1:10,000 dilution) and a alexa488-conjugated goat-anti chick secondary. Other primary antibodies used were mouse anti-RGS14 (Neuromab) (1:300 dilution), rabbit anti- PCP4 (Sigma) (1:600 dilution), guinea pig anti-vGlut2 antibody (Milipore) (1:10,000 dilution), rabbit anti-parvalbumin antibody (Swant) (dilution 1:2000). Alexa-546-conjugated streptavidin (life technologies), secondary antibodies and far-red neurotrace (life technologies) incubations were carried out in block solution for 4 hours at room temperature. Images were collected with a Zeiss 710 laser-scanning confocal microscope.

Reconstructed neurons were classified as either PNs or INs based on the extension and localization of their dendrites and axons. CA1, CA2 and CA3 PNs were identified based on their somatic localization, dendritic arborization and presence of thorny excrescences (TE).

Among INs with somas located in the pyramidal layer (stratum pyramidale, SP), discrimination between BCs and non-BCs was achieved based on the restriction of their axons to SP or not, respectively. When available, firing patterns upon injection of depolarizing current step injection, action potential (AP) half-width, amount of repolarizing sag current upon hyperpolarization from −70 mV to −100 mV by current step injection, membrane resistance (R_M_) and capacitance (C_M_) were additionally used for cell identification. CA2 and CA3a PNs displayed similar firing patterns, AP width, sag current, R_M_ and C_M_. In contrast, INs had faster firing rates, shorter AP width, higher R_M_ and lower C_M_ than PNs. BCs further differed from non-BCs by the presence of a larger sag current. All recorded neurons that could not be unequivocally identified as PNs or INs were excluded from analysis.

### Data analysis and statistics

Electrophysiological recordings were analyzed using IGORpro (Wavemetrics) and Clampfit (Molecular devices) software. For accurate measurements of the kinetics and latencies of post-synaptic responses, the following detection process was used. For each cell, average traces were used to create a template waveform that was then fitted to individual traces and measurements were performed on the fitted trace. When only amplitudes of responses were needed, standard average peak detection was used. Results are reported ± SEM. Statistical significance was assessed using χ^2^ test, Student’s T test, Mann-Whitney U test, Wilcoxon signed-rank test, Kolmogorov-Smirnoff test, Kruskal-Wallis test, one-way or two-way ANOVA where appropriate.

## Author Contributions

RAP, VR & TM designed experiments. RAP, VR, VC, LT, RB, AJYH performed experiments. VR, RAP and DP completed analysis. VR and RAP wrote the manuscript with input from all authors.

## Acknowledgments

This work was supported by the RIKEN Center for Brain Science (TJM), Grant-in-Aid for Scientific Research from MEXT (19H05646; T.J.M), Grant-in-Aid for Scientific Research on Innovative Areas from MEXT (19H05233; T.J.M), ANR-13-JSV4-0002-01 (RAP), ANR-18-CE37-0020-01 (RAP), the Ville de Paris Programme Emergences (RAP), and the Brain and Behavioral Research Foundation NARSAD Young Investigator Grant (RAP) and the Foundation Recherche Médicale, FRM:FTD20170437387 (VR).

## Supplemental figure legends

**Supplemental Figure 1.**
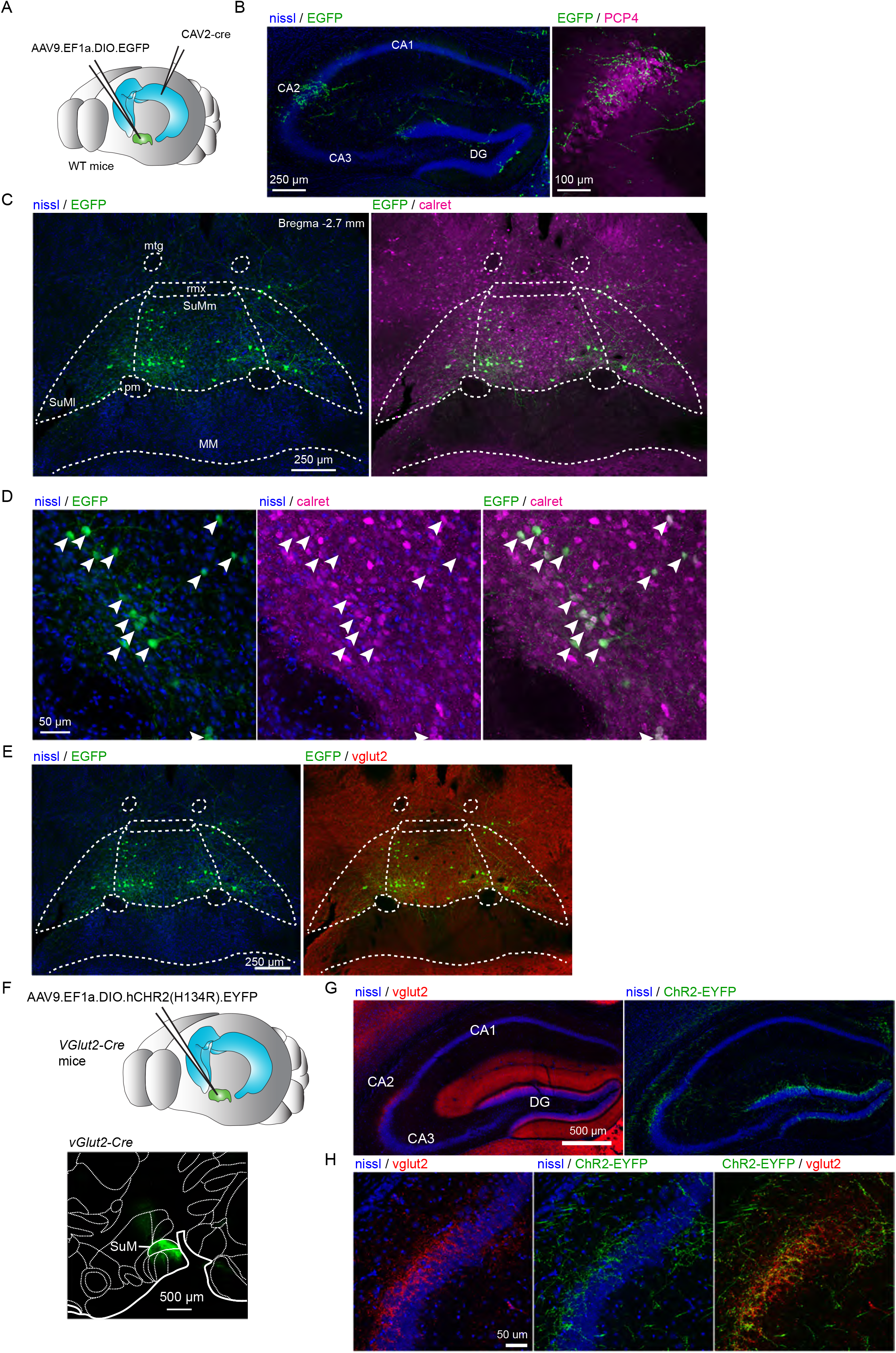
A. Diagram illustrating the intersectional strategy used to label CA2-projecting SuM neurons. B – E. Labelling of CA2-projecting SuM neurons with the retrograde CAV-2 carrying Cre-recombinase injected in CA2 and the anterograde AAV carrying DIO-EGFP injected in SuM of wild type mice. B. Labelling of SuM fibers in the hippocampus from CA2-projecting SuM neurons. Left, nissl staining (blue) and EGFP expression (green) in the hippocampus. Right, PCP4 staining (magenta) and EGFP expression (green) in area CA2. C. Retrograde-labeled SuM neurons that project to hippocampal area CA2. Left, nissl staining (blue) and EGFP expression (green) in SuM. Right, calretinin staining (magenta) and EGFP expression (green) in SuM. D. Higher magnification image of CA2-projecting SuM neurons. Left, nissl staining (blue) and EGFP expression (green) in SuM. Center, nissl (blue) and calretinin staining (magenta) in SuM. Right, calretinin staining (magenta) and EGFP expression (green) in SuM. E. VGluT2 expression of CA2-projecting SuM neurons. Left, nissl staining (blue) and EGFP expression (green) in SuM. Right, VGluT2 staining (red) and EGFP expression (green) in SuM. F. Top, diagram illustrating the injection of AAVs into the SuM. Bottom, sagittal image of the injection site in SuM to express hCHR2(H134R)-EYFP (green) in the VGluT2-Cre line. G and H. Anterograde labelling of SuM projections to the hippocampus from AAV carrying DIO-ChR2-EYFP injected in SuM of VGluT2-Cre mice. G. Left, VGluT2 (red) and nissl staining (blue) in the hippocampus. Right, hCHR2(H134R)-EYFP -expressing SuM fibers (green) and nissl (blue) staining in the hippocampus. H. Left, higher magnification image of area CA2 with VGluT2 (red) and nissl (blue) staining. Center, hCHR2(H134R)-EYFP -expressing SuM fibers (green) and nissl staining (blue). Right, hCHR2(H134R)-EYFP -expressing SuM fibers (green) and VGluT2 staining (red).

**Supplemental Figure 2.**
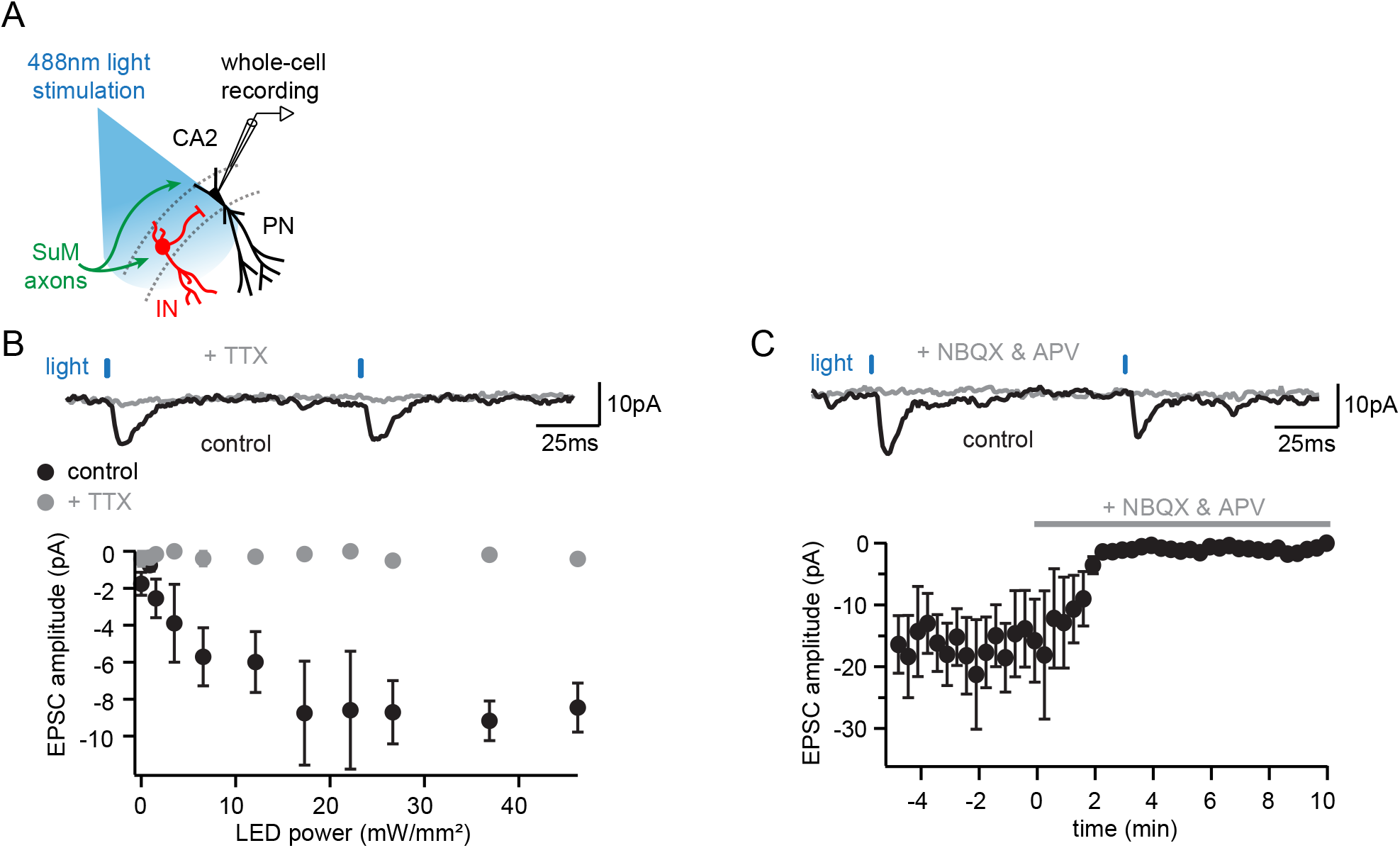
A. Diagram illustrating the whole-cell recording configuration of PNs in area CA2 and SuM fiber stimulation in acute hippocampal slices. B. Light-evoked EPCSs from SuM inputs are completely blocked following application of tetrodotoxin (TTX). Sample traces (top, control shown in black, +TTX shown in grey) and power-response curves (bottom) of light-evoked EPSC amplitudes recorded in PN before (black) and after application of 0.2 μM TTX (grey) at different light intensities (n = 5, error bars represent SEM). C. Light-evoked EPCSs from SuM inputs are completely blocked following application of NMDA and AMPA receptor blockers (NBQX & APV). Sample traces (top, control shown in black, NBQX & APV shown in grey) and time course (bottom) of light-evoked EPSC amplitudes upon application of 10 μM NBQX & 50 μM APV (n = 6, error bars represent SEM).

**Supplemental Figure 3.**
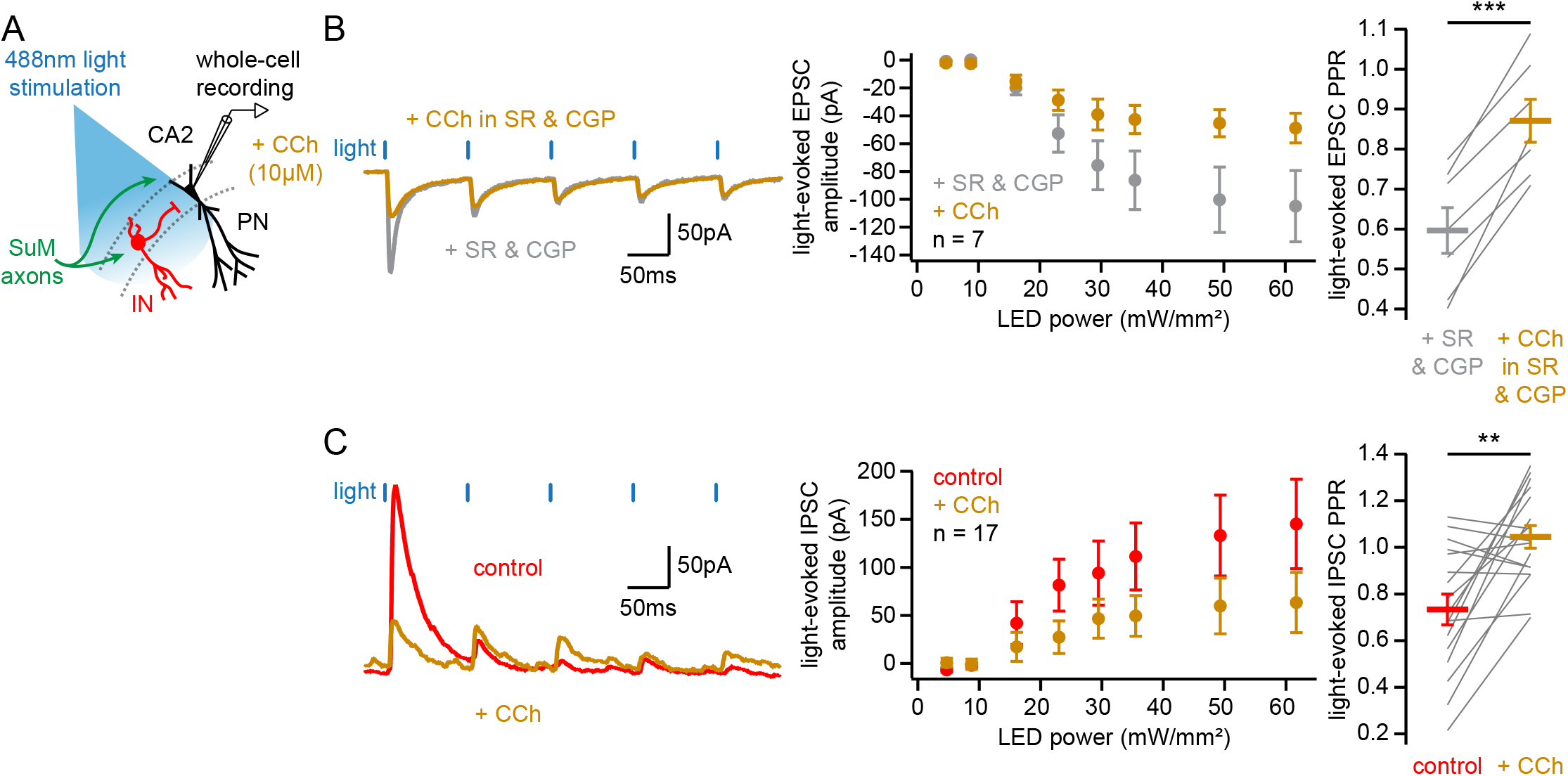
A. Diagram illustrating the whole-cell recording configuration of PNs in area CA2 and SuM fiber stimulation in acute hippocampal slices. B and C. Effect of 10 μM CCh on SuM light-evoked PSCs recorded in CA2 PNs under different conditions : voltage clamp at −70 mV with inhibitory transmission blocked (B, SR95531 & CGP55845A shown in grey, SR95531 & CGP55845A + CCh shown in orange), and voltage clamp at +10 mV (C, control shown in red, CCh shown in orange). Left, sample traces. Middle, power-response curves (B, n = 7; two-way ANOVA with repeated measures, p < 0.001; C, n = 17; two-way ANOVA with repeated measures, p < 0.001; error bars represent SEM). Right, comparison of PPRs (B, n = 7; paired-T test, p < 0.001; C, n = 17; paired-T test, p = 0.001; individual cells shown as grey lines, population average shown as horizontal line, error bars represent SEM).

